# Development of synthetic genetic arrays for studying interactions between the mitochondrial and nuclear genomes

**DOI:** 10.64898/2026.11.16.766915

**Authors:** K. L. Schneider, S. Braam, X. Hao, S. Hanzén, M. Ott, M. Cvijovic, T. Nyström

**Author notes:** Discovery Sciences, AstraZeneca AB R&D, 43183 Mölndal, Sweden. Cochlear Nordic AB, 435 33 Mölnlycke, Sweden.

## Abstract

Mitochondria, the central axis of metabolism and energy transformation has emerged as a focal point for understanding the aging process and a plethora of human diseases. Yet, how the mitochondrial and nuclear genomes communicate is still unclear. This work aims to allow the analysis of such crosstalk by developing a genome-wide platform for studying genetic interactions between the mitochondrial and nuclear genomes. Using yeast as a model system in which mitochondrial homoplasmy and stable alterations in the mitochondrial genome can be generated by biolistic bombardment, we created a genome-wide library of yeast cells carrying nuclear deletions and temperature-sensitive mutations together with a superfolder-GFP reporter for the expression of the mitochondrial-encoded *COX2* gene. Using this approach, we found links between nuclear-encoded processes and mitochondrial gene expression not previously anticipated, including a requirement for genes involved in MVB (multivesicular body) sorting/RIM signaling, vesicle trafficking/glycosylation, polarity, and peroxisome biogenesis, for maintaining mitochondrial *COX2* expression. In contrast, defects in the autophagy machineries, ribosome biogenesis, and nuclear transcription and chromatin organization resulted in increased expression of *COX2*. While reduced *COX2* expression correlated with a reduced expression of the nuclear-encoded *TOM70* gene in mutants defect in mitochondrial functions, expression of the mitochondrial and nuclear genes responded differently to defects in cytoplasmic ribosome biogenesis and autophagy. When creating the *COX2* expression library, we also found that cells devoid of mitochondrial DNA rely on the Rim101 signaling pathway, and many of its auxiliary regulatory factors, to maintain fitness. As mutations in the Rim101 signaling pathway also caused down-regulation of *COX2* expression, it appears that Rim101 signaling is central for maintaining proper control of mitochondrial function. We believe the approach will allow for future genome-wide studies on how nuclear-encoded processes influence the translation, protein folding, and processing of mitochondria-encoded genes.

## Introduction

Mitochondria are membrane-bound organelles responsible for a variety of activities including cellular energy transformation, synthesis of phospholipids, heme, and Krebs cycle intermediates, calcium homeostasis/signaling, and apoptotic activation and cell death. They have retained their own genome, of bacterial evolutionary origin, encoding only a handful of respiratory proteins and mitochondrial tRNAs. Thus, production and assembly of the large composite respiratory chain complex, as well as mitochondrial biogenesis in general, requires a complex and elaborate coordination of nuclear- and mitochondrial-encoded gene expression. Failures to do so due to defects/mutations in mitochondrial DNA (mtDNA) manifest across a spectrum of diseases, including MELAS (Mitochondrial Encephalomyopathy with Lactic Acidosis and Stroke like episodes), MERFF (Myoclonus Epilepsy with Ragged Red Fibres), Leigh syndrome, and Leber’s hereditary optic neuropathy (Copeland 2008). Such diseases, and others resulting from mutations in mtDNA, do not obey Mendelian inheritance as they are inherited through the maternal line only and show unpredictable severity of expression due to heteroplasmy of the mtDNA population (i.e. a mixture of wild type and mutant mtDNA coexisting). Mitochondrial dysfunction, resulting from either mtDNA or nuclear DNA (nDNA) defects, is implicated also in numerous age-related disorders, including neurodegenerative and metabolic diseases, as well as endocrine-related aberrancies and ischemic injury in the heart and brain (Amorim et al., 2022; Fang et al., 2016; Schon et al., 2012). The mitochondrion has also emerged as a key organelle regulating aging itself as well as the performance of other subsystems of the cell, such as the protein quality control in both the cytoplasm and nucleus (Caballero et al., 2011; Higuchi et al., 2013; Hill et al., 2016; Malmgren Hill and Nyström, 2016; Scheckhuber et al., 2007; Suhm et al., 2018b). Moreover, mitochondria have been shown to display an early functional decline during cellular aging (Gottschling and Nyström, 2017; Hughes and Gottschling, 2012), which is suggested to ‘drive’ the aging process, at least in yeast.

While mitochondrial defects can cause aging by a sequential degeneration of interconnected subsystems and organelles (Dillin et al., 2014; Gottschling and Nyström, 2017), some mitochondrial dysfunctions actually extend life span. For example, yeast mitochondrial dysfunction activates intracellular signaling pathways, including the retrograde response and mitochondrial back-signaling (Jazwinski, 2013; Jazwinski and Kriete, 2012), that retards replicative aging through metabolic rewiring and inter-organelle coordination of ribosomal biosynthesis, respectively. Another example in yeast is the demonstration that inactivating members of the mitochondrial translation control (MTC) module required for *COB* and *COX* translation cause a robust boost in sirtuin-dependent genomic silencing and an extension of replicative life span (Caballero et al., 2011). In addition, genetic modifications that block mitochondrial fission delay ageing without impairing fitness in both budding yeast and filamentous fungi (Scheckhuber et al., 2007). Consistent with some mitochondrial dysfunctions triggering lifespan extension in yeast, analogous systems are at work in metazoan, including worms and flies (Copeland et al., 2009; Dell’agnello et al., 2007; Dillin et al., 2002; Feng et al., 2001), in which mutations or RNAi perturbations of the mitochondrial electron transport chain (ETC) subunits have been shown to extend longevity.

Such data makes mitochondria a focal point in understanding cause-and-effects in the aging process. Yet, how mitochondria and the quality control subsystems of other compartments crosstalk and communicate is still unclear. Here, we developed a genome-wide protocol to analyze such crosstalk by creating synthetic gene arrays allowing for the identification of mitochondrial–nuclear genetic interactions in a homoplasmic mitochondrial population. This development allowed us to discover hitherto unknown relationships between nuclear-encoded processes and mitochondrial gene expression.

## Results

### Construction of a mtDNA-less mutant gene array

To construct a yeast collection consisting of single deletions and ts-mutations across the genome that express mitochondrially-encoded reporters, we exploited the power of the synthetic genetic array methodology (SGA, Tong et al., 2001). We created the collection with a two-step crossing protocol, which entails yeast mating and selection for desired genotypes through the growth media and was performed with robotic automation (see Liu et al., 2010). In the first step (Figure 1A), we crossed the single deletion array (xxxΔ, Yeast Knock Out; YKO library) and the essential gene mutant array (temperature-sensitive, ts library) with a strain lacking the gene encoding the mitochondrial DNA polymerase Mip1 (Y7092 *mip1*Δ). Mip1, the only mitochondrial DNA polymerase in yeast, is required for mtDNA replication and is highly conserved and shares approximately 40% amino acid sequence identity with the human polγ catalytic subunit (Lecrenier et al., 1997; Ropp and Copeland, 1996). Yeast cells without Mip1 lose their mtDNA after a few cell divisions, resulting in petite colonies (rho^0^) with cells that fail to perform oxidative phosphorylation and therefore cannot grow on purely respiratory substrates. Consequently, the first crossing step generated a petite version of the mutant arrays, i.e. a YKO and a ts library lacking mitochondrial DNA.

**Figure 1.**
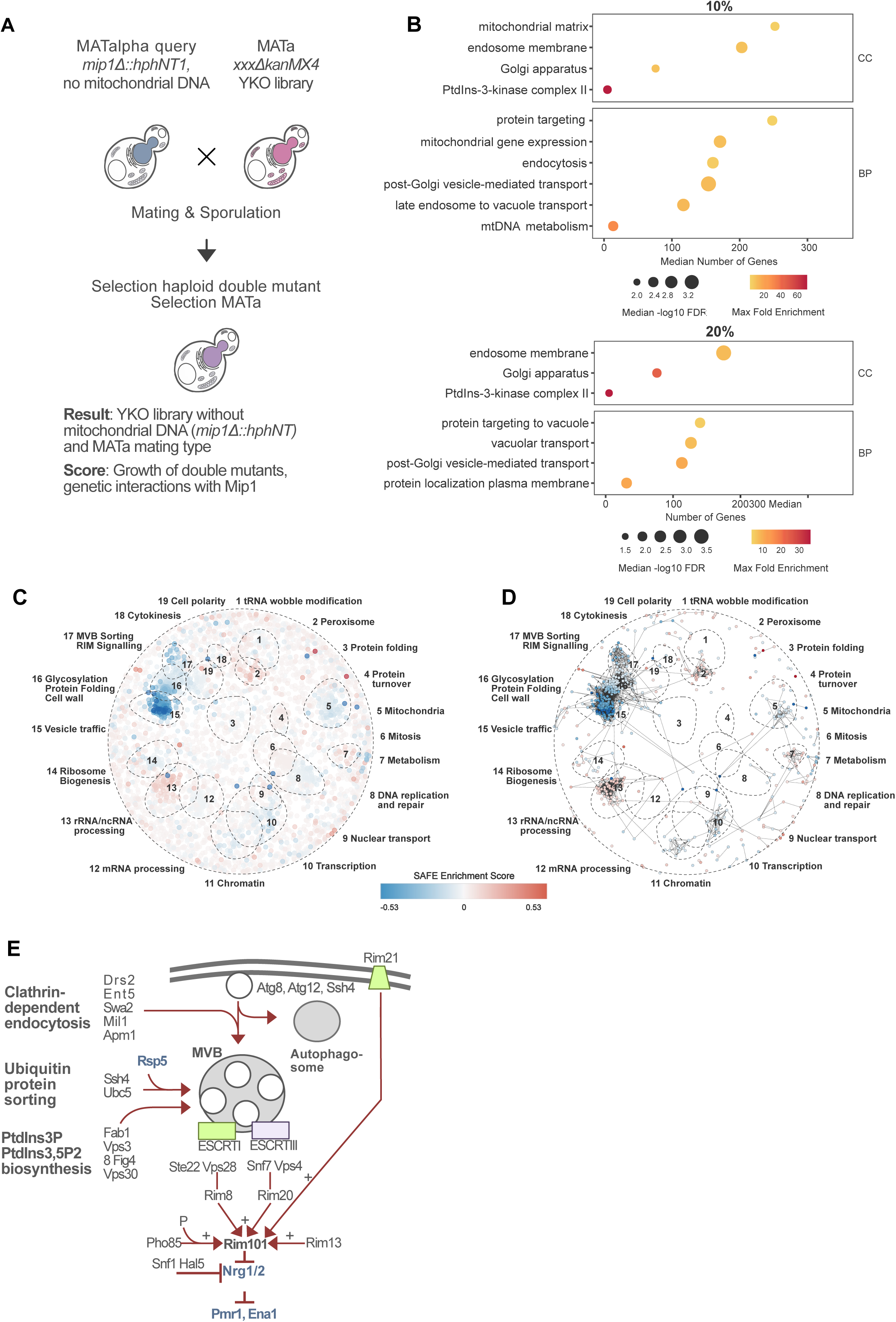
**A:** Strain construction for screening of single gene mutants without mitochondrial DNA by deletion of the mitochondrial DNA polymerase Mip1. **B:** GO-terms of strongest negative genetic interactions of strains without mitochondrial DNA 10% upper panel, 20% lower panel. Using ShinyGo 0.81, generated GO-terms were selected by FDR <0.05 and sorted by fold enrichment. Up to 20 GO-terms were clustered for better readability BP: biological process, CC: cellular component. **C:** Ranked SAFE enrichment landscape across all screened genes. **D**: Subnetwork of genes contributing the strongest to SAFE enrichment, selected from ranked SAFE landscape. Enrichment score >|0.081|. **E:** Detailed illustration of Rim101 pathway. As indicated in SAFE and GO-term analysis, negative genetic interactions upon *MIP1* deletion indicate perturbance of the upstream endosomal trafficking components in the Rim101 pathway, while Rim101 itself and illustrated downstream targets (blue) are not affected.

### Synthetic genetic interactions of single deletion mutants and *mip1Δ*

Due to the *MIP1* homology between yeast and human polγ, yeast Mip1 has been used to study several diseases related to failures in replicating the mitochondrial genome, including Alpers-Huttenlocher syndrome, progeria and peripheral nervous system disease (Copeland, 2008). To investigate the physiological role of the mitochondrial DNA polymerase on a more global level we scored for synthetic genetic interactions between single deletion mutations and the deletion of *MIP1,* using SGA technology (Figure 1A, Supplementary table 1 and 2). For synthetic sick strains, i.e. double *mip1*Δ xxxΔ mutants that grow more poorly (more than a simple additive effect) than the corresponding single mutants, we found gene ontology (GO) enrichments in many processes and components involved in vesicle transport and Golgi function (Figure 1B; 10% and 20% of the mutants with the most severe fitness defect). Spatial Analysis of Functional Enrichment (SAFE) analysis (Baryshnikova, 2016) further pinpointed multivesicular body (MVB) sorting and RIM signaling as processes required for fitness in cells lacking Mip1 (Figure 1C and D). Indeed, we found that the removal of factors of almost the entire *RIM101* signaling pathway were synthetic sick in combination with *mip1*Δ (Figure 1E). RIM signaling is the yeast genetic response to changes in the extracellular pH, ultimately activating the key transcription factor Rim101. This pathway is regulated by ESCRT (endosomal sorting complexes required for transport) complexes, which also play a role in MVB formation, sorting and other cellular functions (Morita, 2012).

We also manually examined the SGA plates for synthetic lethal phenotypes (i.e. no growth of double mutants; Supplementary table 3). We found 16 ORFs across the two runs, of which 15 were synthetic lethal in both runs. Two ORFs were classified as dubious by SGD. The remaining 13 ORFs were analyzed using thecellmap.org (Usaj et al., 2017) with a 0.05 significance cutoff revealed an enrichment in the category mitochondria (Supplementary Figure 1A, supplementary table 5A). The functions of genes lethal in combination with *mip1*Δ span mitochondrial tRNA modifications, mitochondrial translation, and mitochondrial helicase-ssDNA binding- and protease-dependent maintenance of mitochondrial DNA (Supplementary table 3). Regarding positive interactors, we identified 10 deletion mutants in both screens that showed increased growth in the *mip1*Δ background (Supplementary table 4). These positive interactors of mtDNA loss were enriched in the categories ribosome biogenesis, as well as glycosylation and protein folding in thecellmap.org (Supplementary Figure 1B, supplementary table 5B). GO term analysis using ShinyGO did not reveal significant enrichment of functions causing positive genetic interactions with *MIP1*.

### Construction of a genome-wide mutant library with a reporter for mt*COX2* expression

After analyzing *mip1*Δ genetic interactions, we proceeded with constructing a mutant array expressing a mitochondrial genome reporter, *COX2*::sfGFP, together with the wild type *COX2* allele, which encodes a subunit of mitochondrial respiratory chain complex IV (Figure 2A and B). The mitochondrial superfolder GFP (sfGFP), established by Suhm et al., 2018a, serves as a reporter for mt*COX2* expression and was introduced into the mt*COX2* gene by biolistic bombardment (microprojectile bombardment using DNA-coated tungsten particles) as described previously (Suhm et al., 2018a). We inserted the sfGFP into the mitochondrial genome of a Mat-alpha query strain that was subsequently crossed back with the previously created mtDNA-less Mat-a YKO library (Figure 2B) and the ts-mutant library. To ensure the new mutants retained *MIP1*, we additionally selected for *MIP1* with an antibiotic resistance marker (*natNt2*) linked to the *MIP1* gene. The expression of mt*COX2*::sfGFP in the mitochondrial genome is facilitated by *COX2* UTRs but under standard growth conditions with glucose as the carbon source, mitochondrial genome expression is very low. Therefore, we performed the high content-microscopy screen using glycerol instead of glucose as the carbon source. Mutant strains unable to grow in glycerol, such as those with disrupted glycerol uptake or metabolism, would therefore be lost during cultivation prior to imaging. We also tagged the nuclear-encoded mitochondrial importer Tom70 with mScarlet-I (Schneider et al., 2021) to be able to visualize mitochondria independently of mt*COX2*::sfGFP expression and to allow for a comparison between mt*COX2*::sfGFP expression and mitochondrial content, number, and morphology. The Tom70-mScarlet-I signal was furthermore used as a mitochondrial mask to verify mitochondrial expression of mt*COX2*::sfGFP (Figure 2C and D). Figure 2E shows an example of strains displaying lower and higher mt-sfGFP signals compared to wildtype and the Tom70-mScarlet-I signal.

**Figure 2.**
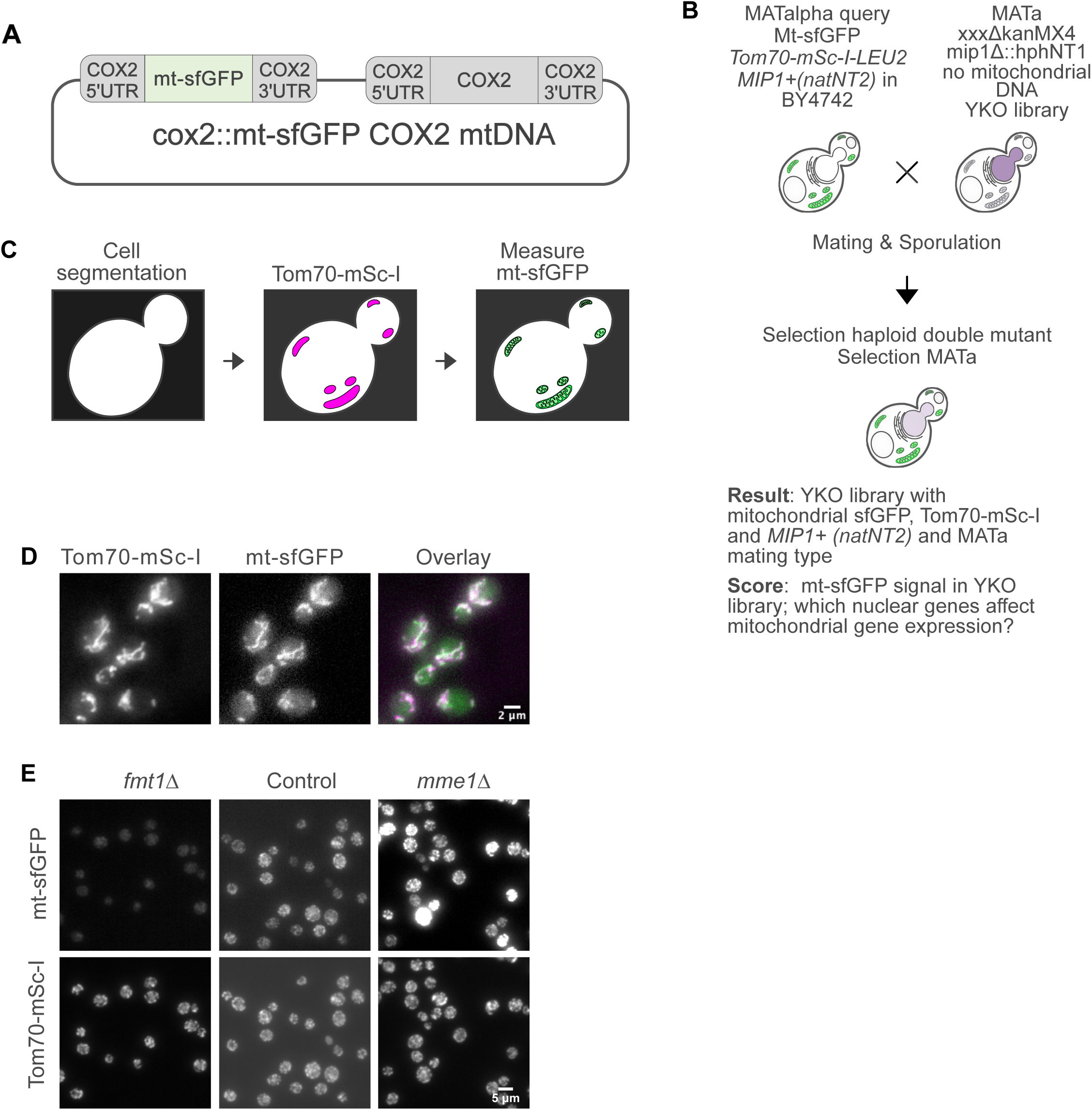
**A:** Schematic of integration of sf-GFP cassette into *COX2* locus within mitochondrial DNA. **B:** Strain construction for screening of single gene mutants with mitochondrial sfGFP expression and Tom70-mSc-I in homoplasmic strains containing mitochondrial DNA with the described sfGFP expression casette. **C:** Illustration of automated image analysis algorithm. Cell segmentation was followed by creating a mask for the localization of mitochondria to measure mt-sfGFP intensity. **D:** Exemplary images generated during high content microscopy, scale bar: 2 μm. **E:** Representative images of control strain, and *fmt1Δ* and *mme1Δ* strains as example for cells with higher or lower sfGFP signal, respectively. Scale bar: 5 μm.

### Validation of the screening procedure and analysis of global relationships between *COX2* expression and mitochondrial content, number, and morphology

Cells were imaged in 96-well plates using a high content-microscope and we observed varying normally distributed mt*COX2*::sfGFP expression, as expected (Figure 3A and B; Supplementary Figure 2A-D). It should be noted that we use the term expression here in the original and broad sense without linking it to a particular process; in other words, altered expression could be due to altered transcription, translation, mtDNA copy number, altered protein stability etc. However, we could normalize expression to total mitochondrial content and/or mitochondrial number using the Tom70-mScarlet-I signal and visual inspection revealed that the GFP signal consistently overlapped with the Tom70-mScarlet-I signal. We employed the MetaXpress software to overlay the Cox2-sfGFP signal with mitochondrial masks generated in the RFP channel and determined mean GFP fluorescence intensity (arbitrary unit) within the masks as the primary readout for this screen. We extracted Cox2-sfGFP fluorescence intensity per cell for each well with standard deviation for these values. In addition, we collected data on mitochondrial area, number of connected/tubular mitochondria per cell, and RFP intensity, all determined by the masks generated in the Tom70-mScarlet-I channel. Validating the control values for these three parameters for each plate provide a reliable reference point (Figure 3C-H). Most plates show a normal distribution and the weighted mean of control values per plate coincides closely with the peak of the distribution. In some plates control values were slightly skewed, typically limited to one parameter as illustrated by the mScarlet-I intensity density plot for plate 10 (Supplementary Figure 2B, plate 10). However, no technical issues, such as poor focus during imaging, were found for this plate, suggesting that the somewhat skewed values might accurately reflect the data, indicating unexpectedly low mScarlet-I intensity values for the mutants on this particular plate. Given that the SGA deletion array is ordered by chromosomal gene location, pathway-related genes may cluster on a single plate and their co-regulation could plausibly account for the observed skew.

**Figure 3:**
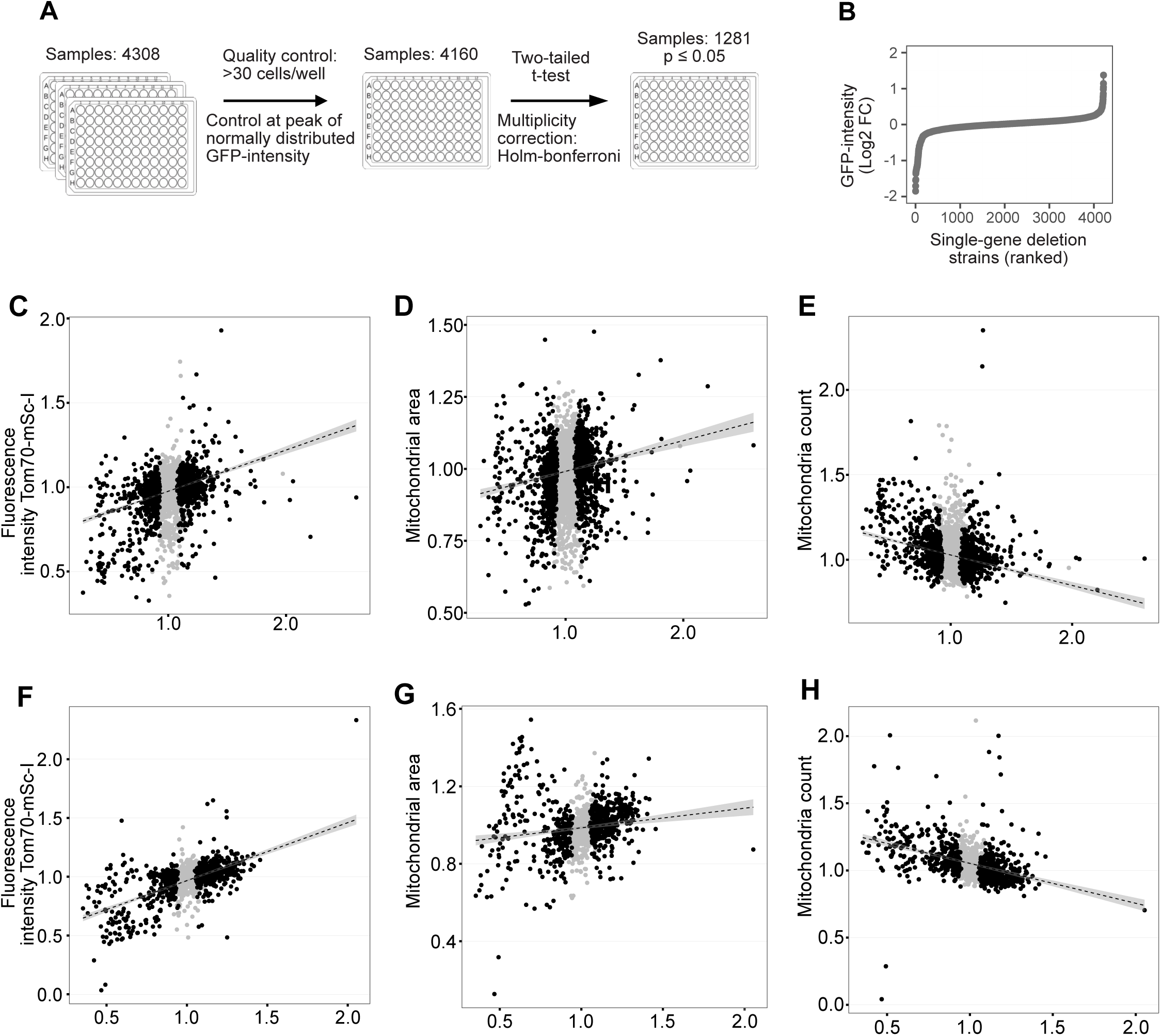
**A:** Process of quality control and t-testing of fluorescence intensity detected during high content microscopy. Control strains were coerced by calculating weighed mean per plate and t-testing was done per plate to account for in-plate variation**. B:** Distribution of measured sfGFP intensity across the single-gene deletion library. Extreme outliers were excluded to ensure readability (total of 7). **C-E**: Correlation between GFP intensity and various measured mitochondria characteristics, all displayed as plate-wise fold change from control strain. Strains expressing sfGFP derived from single gene deletion array. **F-H:** Display of correlations for temperature sensitive strains. All graphs show the measured platewise raw fold change relative to wildtype of indicated mitochondrial characteristic measured over GFP intensity if not labeled otherwise. Black dots: adjusted significant p-value, grey: non significant adjusted p-values. Dashed line and grey shading: linear regresssion of all datapoints (significant and non-significant p-value).

Using the GFP intensity values, we performed a t-test to identify mutants significantly differing from the weighted mean control values per plate. Comparing the test strains to control strains on each plate individually helped correct for differences in handling and imaging between plates and days. Density plots of these values across the entire YKO collection indicated a normal distribution (Supplementary Figure 2E, left). This distribution became more pronounced when outliers were removed (Supplementary Figure 2E, center). However, we decided to include outliers in our downstream analyses, as they represent extreme values with both higher and lower mt-sfGFP expression (Supplementary Figure 2E, right). We suspect these mutant strains to be actual hits in our screen as visual inspection confirmed successful image acquisition and noticeable deviations of mt-sfGFP compared to control strain images.

Plotting the GFP intensity measurements for each mutant strain revealed three populations of hits (Figure 3C-E). As expected, most values clustered around the control. A few hits had a higher ratio of mtCox2-sfGFP/control than the wild type strains, while a larger population of strains had a lower ratio. This indicates that there are more mutants in the YKO collection that exhibit reduced mtCox2-sfGFP levels than those that exhibit increased levels, but both populations of mutants exist across the library. These three populations were observed by plotting the mtCox2-sfGFP intensity values against readouts obtained through the Tom70-mScarlet-I signal; i.e. RFP intensity (Figure 3C), mitochondrial area (Figure 3D), and mitochondria count (Figure 3E). Using these values, we focused on detecting potential general patterns of correlations between *COX2* expression and mitochondrial content/morphology. We performed linear regression of the mtCox2-sfGFP values plotted against Tom70-mScarlet-I data. We found a general positive trend between mtCox2-sfGFP levels and Tom70-mScarlet-I intensity and mitochondrial area (Figure 3C and D). These data suggest that increased mitochondrial content/biomass in the cell often correlates with increased mtCox2-sfGFP intensity and that the expression of the mtDNA-encoded *COX2* and the nDNA-encoded *TOM70* are most often responding similarly to any given mutation. The mitochondrial count, used to indicate mitochondrial fragmentation, decreased with higher mtCox2-sfGFP intensity indicating a general decreased *COX2* expression with mitochondrial fragmentation (Figure 3E).

We performed the same statistical analysis for the temperature-sensitive mutant library of essential genes. The linear regression analysis indicated that the trends between mtCox2-sfGFP levels and mitochondrial count, area, and Tom70 levels we observed in the YKO collection analysis were consistent also for essential gene mutants (Figure 3F-H). Additionally, both libraries had an apparently random distribution of hits across the yeast chromosomes (Supplementary Figure 2F and 3F) and the density plots of mtCox2-sfGFP levels indicated normal distribution (Supplementary Figure 2A and 3A). The outliers, including extreme values, within the temperature-sensitive library were especially frequent for decreased mtCox2-sfGFP levels compared to the control (Supplementary Figure 3E, right panel).

The temperature-sensitive collection contains many mutant alleles of the same genes for 30% of the collection, including 20 alleles of *ACT1*, 6 alleles of *POL1*, 7 of *CDC24* (Li et al., 2011). Plotting the different alleles available for each ORF according to their fold change in mtCox2-sfGFP level revealed that mutations affecting the same gene (Supplementary Figure 3G), for example *BBP1*, *CDC10*, *GAB1* and *BRL1* can affect mitochondrial gene expression differently. This may not be surprising as temperature-sensitive mutant alleles of the same gene are known to exhibit distinct phenotypes and such differential effects of different alleles may be useful in analyzing the more specific mechanisms affecting mt gene expression.

### Enrichment analysis of mutations causing altered expression of *COX2*

When scoring the mutants displaying reduced *COX2* expression using Spatial Analysis of Functional Enrichment (SAFE, Baryshnikova, 2016), we found, as expected, a strong enrichment for genes involved in mitochondrial functions (Figure 4A and B), showing that mutations in any of a large number of nuclear genes encoding mitochondrial proteins will cause a reduced expression of a mitochondrially-encoded one. Moreover, SAFE analysis identified additional processes affecting *COX2* expression negatively when mutated; including peroxisome biogenesis, cell polarity, glycosylation, vesicle traffic, mRNA processing, and MVB sorting/RIM signaling (Figure 4A). The latter process included the hits *rim101Δ* and *rim20Δ*, highlighting, again, a potential connection between mitochondrial function and RIM signaling (see also Figure 1E). GO term enrichment for the top (with respect to degree of down-regulation) 20% of the hits also identified amino acid and amine biosynthesis and metabolism as processes required for maintaining *COX2* expression (Figure 4C). Similarly, when analyzing whether any of the top 20% hits for decreasing *COX2* expression were part of known protein-protein and/or gene-gene interaction networks using STRING analysis we identified the Lys2-Ser2-Spe1-Mct1 network involved in amino acid, spermidine and fatty acid biosynthesis; some of the proteins in this network are localized to mitochondria (Figure 4E).

**Figure 4:**
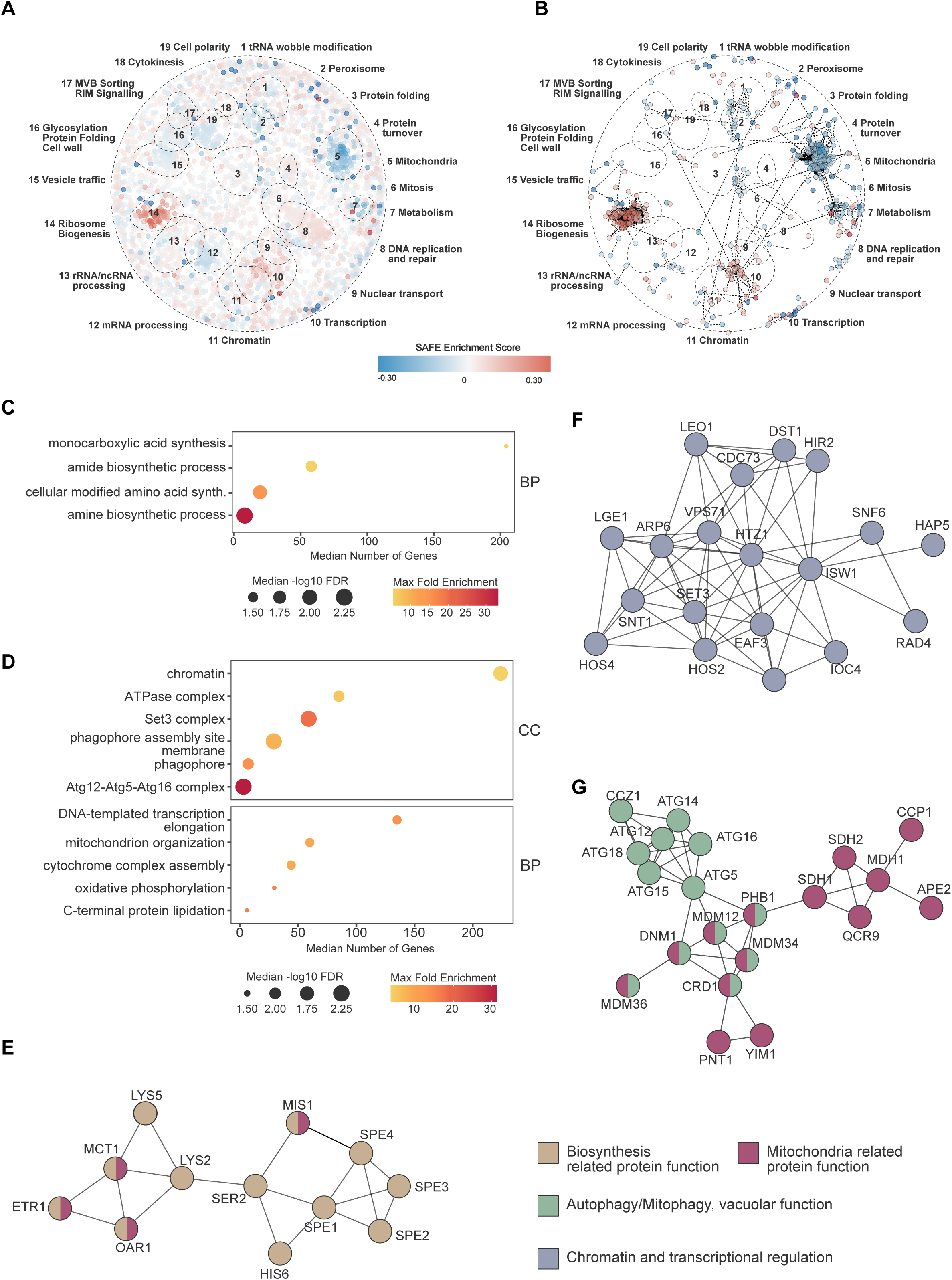
Gene enrichment analyses based on foldchange to control of sfGFP expression in high content microscopy analysis **A:** Ranked SAFE enrichment landscape based on foldchange across all screened genes after quality control. Thresholding has been omitted to show overall global distribution of enrichment. **B**: Subnetwork of genes contributing the strongest to SAFE enrichment, selected from ranked SAFE landscape. Threshold for selection of genes: enrichment score >|0.081|. **C:** Bubble plot of clustered GO-terms for lowest 20% subset of foldchange sfGFP expression compared to wildtype. Generated GO-terms were selected by FDR < 0.05 and sorted by fold enrichment and subjected to semantic clustering to reduce redundancy. Genes in pathway were restricted to ≤ 350. **D:** GO-term enrichment ot top 20% subset of upregulated sfGFP expressing single gene deletion mutants. **E-G:** Protein interaction network generated from 20% lowest (E) or highest (F-G) sfGFP-expressing single-gene deletion strains. Identification of Genes is based on Gene Ontology annotations and edges represent known or predicted protein–protein associations from the STRING database.

SAFE analysis identified three major groups increasing *COX2* expression when mutated; ribosome biogenesis, DNA replication/repair, and transcription (Figure 4A and B). SAFE together with GO term enrichment for the top (with respect to degree of up-regulation) 20% of the hits pinpointed the chromatin-associated histone deacetylase (HDAC) Set3 complex as a major component of the transcriptional apparatus affecting mitochondrial *COX2* expression (Figure 4D). Accordingly, STRING analysis (Szklarczyk et al., 2023), revealed that a large number of the genes involved in chromatin remodeling and transcription found to affect *COX2* expression are inter-connected by both physical and genetic interactions (Figure 4F); these genes include those of the Set3, Htz1, NuA4, and Swr1 complexes.

GO term enrichment also highlighted that mutations in components and processes related to autophagy and mitophagy, including genes encoding components of the Atg12-Atg5-Atg16 complex, the phagophore, and the phagophore assembly site membrane, caused an increased expression of *COX2* (Figure 4D). Additional genes/proteins interacting genetically and/or physically with genes/proteins of the Atg12-Atg5-Atg16 complex identified in the screen included Atg15, required for lysis of autophagic bodies, Atg32, a mitochondrial outer membrane protein required to initiate mitophagy, Ccz1, of the Mon1-Ccz1 GEF complex required for both macroautophagy and mitophagy, Atg14, required to target complex I to the phagophore assembly site (PAS), and Atg9, mediating autophagosomal membrane expansion (Figure 4G).

### Enrichment analysis of mutations causing altered expression of *TOM70*

Since the mutant libraries created also contained the Tom70-mScarlet-I reporter, we were able to determine whether the mitochondrial proteins, Cox2 and Tom70, encoded by the mitochondrial and nuclear genomes, respectively, behaved similarly or differently as a consequence of defects in different specific biological processes. Similar to the expression of *COX2*::sfGFP, *TOM70*::mScarlet-I expression was reduced by mutations in a large number of genes encoding mitochondrial proteins/functions and enhanced by mutations in many genes involved in nuclear transcription and chromatin organization (Figure 5A). In contrast, mutations in genes involved in ribosome biogenesis and translation caused a reduced *TOM70*::mScarlet-I expression as opposed to the case for *COX2*::sfGFP (compare Figures 4A, B and 5A, B). GO term enrichment for the top (with respect to degree of up-regulation of *TOM70*::mScarlet-I) 20% of the hits pinpointed the chromatin-associated histone deacetylase (HDAC) Set3 complex as a component also affecting expression of *TOM70*::mScarlet-I (Figure 5C). In contrast to the case for *COX2*::sfGFP expression, mutations in genes involved in autophagy and mitophagy were not enriched among the genes enhancing *TOM70*::mScarlet-I expression (Figure 5C). The data show that the mitochondrial- and nuclear-encoded genes analyzed respond similarly to the disruption of some specific components and biological processes while acting differently to the disturbance of others.

**Figure 5:**
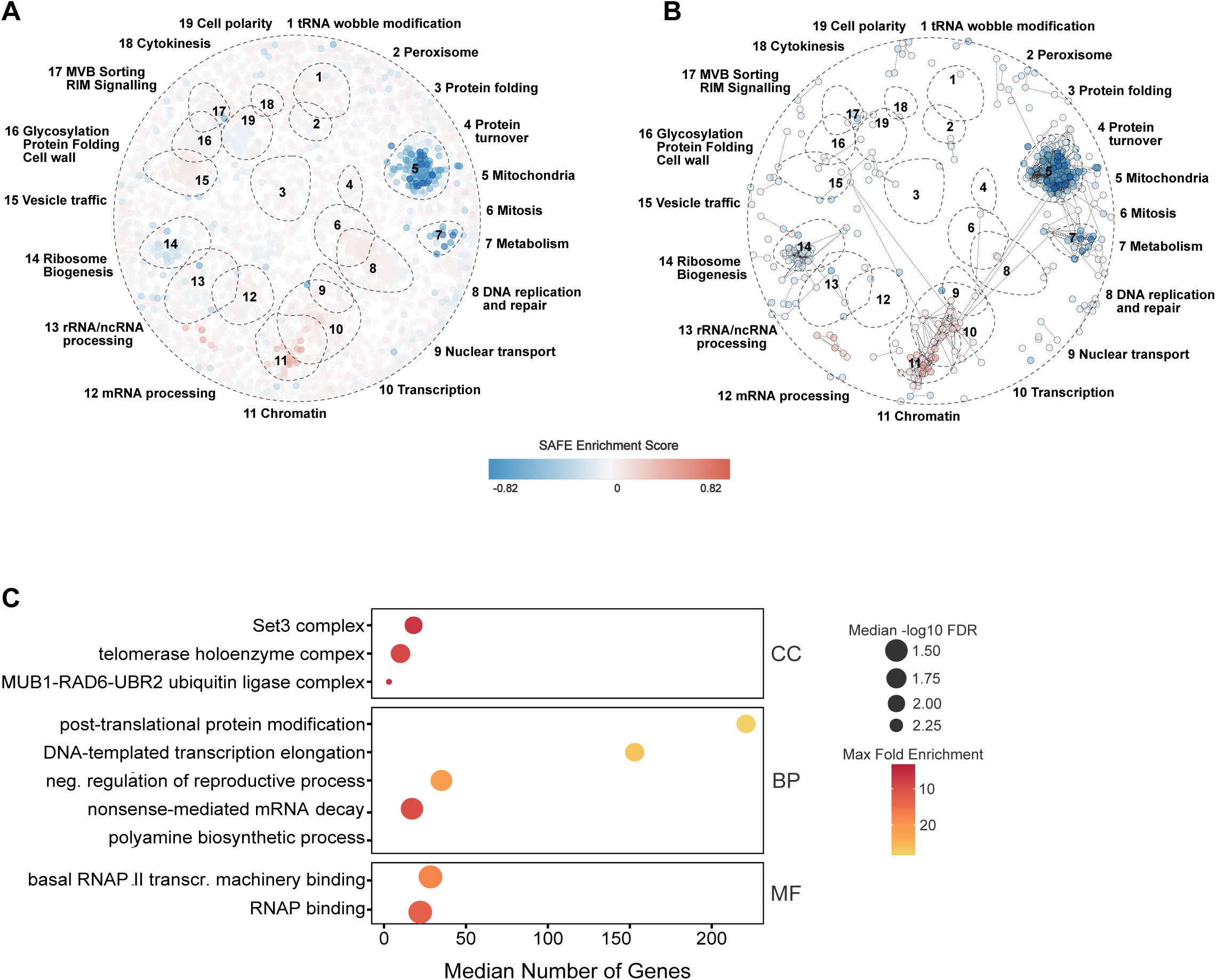
Gene enrichment analyses based on foldchange to control of Tom70 expression in high content microscopy analysis **A:** Ranked SAFE enrichment landscape based on flodchange across all screened genes after quality control. Thresholding has been omitted to show overall global distribution of enrichment. **B**: Subnetwork of genes contributing the strongest to SAFE enrichment, selected from ranked SAFE landscape. Threshold for selection of genes: enrichment score >|0.081|. **C:** Bubble plot of clustered GO-terms for lowest 20% subset of foldchange Tom70 expression compared to wildtype. Generated GO-terms were selected by FDR <0.05 and sorted by fold enrichment and subjected to semantic clustering to reduce redundancy. Genes in pathway ere were restricted to ≤ 350.

## Discussion

In this study, we developed a protocol based on synthetic genetic array (SGA) technology and biolistic transformation of the mitochondrial genome, enabling genome-wide investigation of interactions between the nuclear and mitochondrial genomes. Statistical analyses of the screens confirmed that the method is successful in identifying significant deviations of mtCox2-sfGFP levels in genome-wide mutant libraries and, as expected, expression of *COX2* was reduced by many mutations in the nuclear genes encoding mitochondrial proteins, including positive regulators of mitochondrial translation. However, there are some limitations to our screen setup: The yeast mutant libraries and SGA methodology are usually performed using the S288C genetic background, which display a certain frequency of spontaneous petite formation (Baruffini et al., 2007), but this is also true for many other yeast genetic backgrounds and is difficult to circumvent. Additionally, monitoring mt-sfGFP expression is best performed when culturing cells in glycerol and, therefore, mutants of the glycerol pathways and respiration-deficient mutants will be excluded in the analysis. Possibly, a modified reporter fusion giving a stronger signal from *COX* (or *COB*) might allow for the analysis of expression also on non-respiratory substrates. Furthermore, deletion mutants unable to grow without mtDNA due to the *MIP1* deletion obviously will not survive to the second crossing in the screen generating the mt*COX2*::GFP reporter library. As reported herein, we found 16 such mutants in the library (Supplementary table 3). The function of some of these mutated genes overlap with those of two previous studies (Dunn et al., 2006; Stenger et al., 2020) aimed at finding genes required for yeast cells to grow without mtDNA. However, the exact identity of the genes differs greatly in all three studies. A reason for this discrepancy might be differences in strain backgrounds and experimental protocols as pointed out previously (Stenger et al., 2020); the previous two studies used ethidium bromide to generate mtDNA-less cells (Dunn et al., 2006; Stenger et al., 2020), whereas we removed the mtDNA polymerase. Thus, we suggest that the mutations identified in this work are, at this time, best described as synthetic lethal together with a *MIP1* deletion rather than being mutations in genes that in general are required for yeast to grow without mtDNA. Another limitation of this work, of course, is that the alterations in gene expression observed in the different mutants identified could be due to various factors, such as altered transcription, translation, mtDNA content, and/or mt-sfGFP degradation. Thus, the exact mechanism behind the different levels of mtCox2-GFP observed must be elucidated by follow-up experiments. Nevertheless, we believe that our method may be instrumental in deciphering new pathways affecting mitochondrial gene expression and mitochondrial quality control as the approach and development allow for scoring the entire complement of nuclear-encoded genes affecting gene expression of mitochondrially-encoded genes and, using other reporters, the proofreading and folding of their products.

The protocol and libraries used allowed us to find links between nuclear-encoded processes and mitochondrial gene expression not previously anticipated, including a role for peroxisome biogenesis, cell polarity, glycosylation, vesicle traffic, mRNA processing, and MVB sorting/RIM signalling in sustaining mt*COX2* expression. Notably, the MVB sorting/RIM signalling pathway was required also for sustaining fitness of cells lacking mtDNA due to the *MIP1* deletion. As for nuclear mutations causing an increased mt*COX2* expression, most were found in the GO group of ‘chromatin remodeling/regulation, transcription regulation, histone remodeling’, and especially mutations in genes/proteins of the Set3 histone deacetylase complex and genes/proteins of the Swr1 complex. The latter exchanges histone variant H2A.Z for chromatin-bound histone H2A, and has been shown to be an aging factor that mediates lifespan extension by dietary restriction (Garay et al., 2014). In addition, we found that mutations in the autophagy machinery, another system required for lifespan extension by dietary restriction (Hansen et al., 2008; Taylor and Dillin, 2011), resulted in an increased expression of mt*COX2*. Autophagy and mitochondria are directly linked through the selective elimination of mitochondria, called mitophagy. It is possible that disrupting such a process elevates mitochondrial content and, concomitantly, mtCox2-GFP levels. However, elevated mtCox2-GFP levels in mutants with deletions in autophagy genes was not matched by elevated levels of mtTom70-mScarlet-I, suggesting that the mitochondrial content was not elevated, or at least that the expression of mt*COX2*::GFP and mt*TOM70*::mScarlet-I was not coordinately up-regulated in these mutants. Whether or not the Swr1 complex and autophagy affect lifespan via their effect on mitochondrial gene expression remains to be elucidated.

One take-home message of this work is that the expression of a mitochondrial-encoded and a nuclear-encoded gene, both producing a mitochondrial protein, may, or may not be coordinated. For example, we found a strong coordinated response of the (mt)gene and the (n)gene to mutations in a large number of genes involved in mitochondrial biogenesis and function. In contrast, the (mt)gene and (n)gene responded differently to mutations in ribosomal genes and genes otherwise involved in translation. Such results beg the question of how these defects in nuclear-encoded processes communicate and signal with mitochondrially-encoded and nuclear-encoded expression and if new organelle cross-communication pathways can be discovered based on these findings. We believe the development of our arrays for studying genetic interactions between nuclear- and mitochondrial-encoded genes may help answer such questions.

## Acknowledgments

The authors would like to thank Andreas Kohler, Tamara Suhm and Charles Boone for sharing strains and plasmids. TN acknowledges support of Knut och Alice Wallenbergs Stiftelse (KAW2023.0277 Scholar) and Vetenskapsrådet (VR2023-03763) and MC acknowledges support of Swedish Research Council (VR2023-04319).

## Materials and methods

### SGA technology preparation of screen libraries: SGA screen with *mip1Δ*

#### *mip1Δ* single gene knock-out (YKO) library and *mip1Δ* essential gene (temperature-sensitive) library generation

The libraries were constructed using the SGA technology (Tong et al., 2004; Tong et al., 2001) to introduce *mip1Δ* into the yeast single gene knock-out collection (YKO, SGAV2), and essential gene temperature-sensitive allele collection (ts-V6). The query strain lacking the mitochondrial DNA polymerase Mip1 (Y7092 MAT alpha *his3Δ1 can1Δ::STE2pr-Sp_his5 lyp1Δ ura3Δ0 leu2Δ0 met15Δ0 LYS2+mip1::hphNT1*) was mated with the SGAV2 and ts collections. Y8835 (Y8835 MAT alpha *can1Δ::STE2 pr-his5 lyp1Δ ura3Δ::hphNT1 leu2Δ0 his3Δ1 met15Δ0 cyh2*) was used as the control strain with wild-type *MIP1*. The genotypes are also shown in Supplementary Table 1. The SGAV2 and ts-V6 collections were kindly provided by Prof Charlie Boone, University of Toronto, Canada. Diploids were selected using YEPD+ HygB (Formedium, CAS No. 31282-04-9) + G418 (Fisher Scientific CAS No. 108321-42-2). Then cells were transferred onto sporulation plates for 14 days. After that haploids were selected using Sc-his-arg-lys + Canavanine (L-Canavanine, Sigma Aldrich, CAS No.543-38-4) + S-AEC (S-(2-Aminoethyl)-L-cysteine hydrochloride, Sigma Aldrich, CAS No.4099-35-8). For the final selection Sc-his-arg-lys + Canavanine + S-AEC+HygB+G418 were used. The final selection plates of *mip1Δ* single deletion library were grown at 30°C and imaged after 2 and 3 days. The *mip1Δ* ts library was grown at 22°C /26°C / 30°C and imaged after 2 and 3 days.

#### mt-sfGFP mutant library construction

In the second step, we used the mutant collections without mtDNA we created previously and crossed them with a new query strain according to SGA technology. This query contained mitochondrial DNA expressing markerless *COX2::sfGFP* construct and was constructed by biolistic transformation of BY4742 (expressing TOM70-mSc-I-LEU and a NAT cassette close to the *MIP1* locus) as described previously (Suhm et al., 2018a) (Supplementary table 1). Diploids were selected using Sc-leu+ clonNAT (Werner BioAgents, CAS 96736-11-7) + G418. Then, cells were transferred onto sporulation plates for 14 days. After that haploids were selected using Sc-his-arg-lys-leu + Canavanine + S-AEC. For the final selection Sc-his-arg-lys-leu + Canavanine + S-AEC+G418+ clonNAT were used.

### Mitochondrial stain and imaging

Cells were grown until mid-exponential phase (OD600∼0.5) in YPG (3% glycerol) and mitochondria were stained using rhodamine B hexyl ester perchlorate (Thermo Scientific) at 100 nM final concentration as described previously (Babazadeh et al., 2019). Light microscopy was performed with a conventional fluorescence microscope, the Zeiss Axio Observer .Z1 inverted microscope equipped with an Axiocam 506m camera and a Plan-Apochromat 100x/1.4 NA Oil DIC M27 objective. A representative image is shown in maximum Z-projection of relevant Z-steps. Linear brightness and contrast adjustments were made to highlight co-localization. Figure 2E images stem directly from the high content-imaging screen and were adjusted in a linear manner to the same pixel values across the three images.

### High content-imaging

Cells were grown in complete synthetic media and low-fluorescence yeast nitrogen base (LoFlo, Formedium) supplemented with 3% glycerol. YKO collection strains were grown at 30°C, temperature-sensitive collection strains at 22°C and shifted to 30°C for 2 hours prior to imaging, slow-growing collection strains at 22°C. Pre-cultures were started by thawing 96-well plates from -80°C at RT and adding glycerol stock to media. Plates were incubated shaking for 2 days. Cells were re-diluted in fresh media for growth for 24 hours to OD_600_∼0.5 and harvested in 96-well glass bottom wells by centrifuging 1:1 cell suspension and fresh pre-warmed media. Imaging was performed with live cells using the ImageXpress Micro XLS system (Molecular devices) equipped with a 100X objective (CFI L Plan EPI cc 0 mm to 0.7 mm). A single slice was acquired for up to four fields of view per well. Segmentation and image analysis was done using MetaXpress software (version 6.54.532; Molecular Devices). Cells were counted, mitochondrial masks were generated in the Tom70-mScarlet-I images and mt-sfGFP intensity was measured within the mitochondrial masks. Additionally, Tom70-mScarlet-I images were used to quantify mitochondrial area, count and Tom70-mScarlet-I intensity. It is possible that some of the strains with severe growth defects on nonfermentable carbon sources grew just enough to reach the cutoff cell number in the imaging screen or that such petite mutants in the collections accumulated suppressors. Therefore, it is crucial to manually confirm specific hits of interest, especially those with extreme values. We also inspected images for the two hits with the highest mitochondria count and found that, while the mitochondria appear somewhat fragmented, the values might be overestimated by the segmentation software due to cells clumping together, causing them to be partially out of focus instead of in one Z-layer. Although rare, such occurrences can generate extreme values. Overall, the mt-sfGFP analysis in MetaXpress was effective. We manually inspected the images for the top 20% of the hits with decreased mt-sfGFP signal compared to control (123 mutants) and confirmed focused images without artifacts and cells with visibly low mt-sfGFP signal.

### Screen analysis and statistical testing

If not specified otherwise, statistical analysis and data handling was done in R (version 4.3.2) using the tidyverse collection of packages (R Core Team, 2021; Wickham et al., 2019).

Wells with cell counts below 30 have been removed from analysis, reducing the dataset from 4239 individual genes to 4148 (SGA screen) and from 885 genes (1232 test cases including alleles) to 840 genes (1130 test cases). Platewise, weighted mean and weighted variance were calculated for the control strain to be used in a following two tailed t-test to account for varying cell numbers per well and plate.

Outliers and extreme outliers were determined according to interquartile range criterion (IQC, 1.5 x IQC and 3 x IQC, respectively). GO terms indicated that outliers are found mainly in mitochondrial related function and might represent the actual measurement of the assay, hence all outliers were included in the t-test. Multiple testing was corrected by applying Holm-Bonferroni adjustment. If not indicated otherwise, all measurements with adjusted p-value ≤ 0.05 were considered significant.

Gene Ontology (GO) enrichment term analysis was done with the web interface of ShinyGo (version 0.81, Ge et al., 2020). Generated GO terms were selected by FDR < 0.05 and sorted by fold enrichment score and ontology. Up to 20 terms of the significantly enriched GoTerms with the highest fold enrichment were subjected to semantic clustering using rrvgo (similarity threshold 0.7, minimal cluster size 3) to avoid redundancy for visualization (Sayols, 2023).

STRING protein association network was generated with R package Stringdb on the STRING database version 12.0 with a combined confidence score of >400, corresponding to medium confidence (Szklarczyk et al., 2023).

SAFE analysis was performed and visualized using the Cytoscape SAFE app (Cytoscape version 3.10.4, Assenov et al., 2008; Baryshnikova, 2016; Morris et al., 2014; Raudvere et al., 2019; Shannon et al., 2003). Nodes were mapped as log2 transformed fold change to wildtype on the genetic interaction similarity network from Constanzo et al. (2016) and significance of neighborhood enrichment was assessed with SAFE permutation based statistical testing as described in Baryshnikova 2016.

## Notes

### Competing Interest Statement

KLS is currently employed by AstraZeneca AB and SH is currently employed by Cochlear Nordic AB. All experimental research reported in this study was performed at the University of Gothenburg before the authors joined their respective companies. Neither company participated in the experimental research or influenced the study design, data collection or interpretation of the results. The authors declare no other competing interests.

