## Supplemental figure 1 for "Development of synthetic genetic arrays for studying interactions between the mitochondrial and nuclear genomes"

### Supplementary Figure 1

A

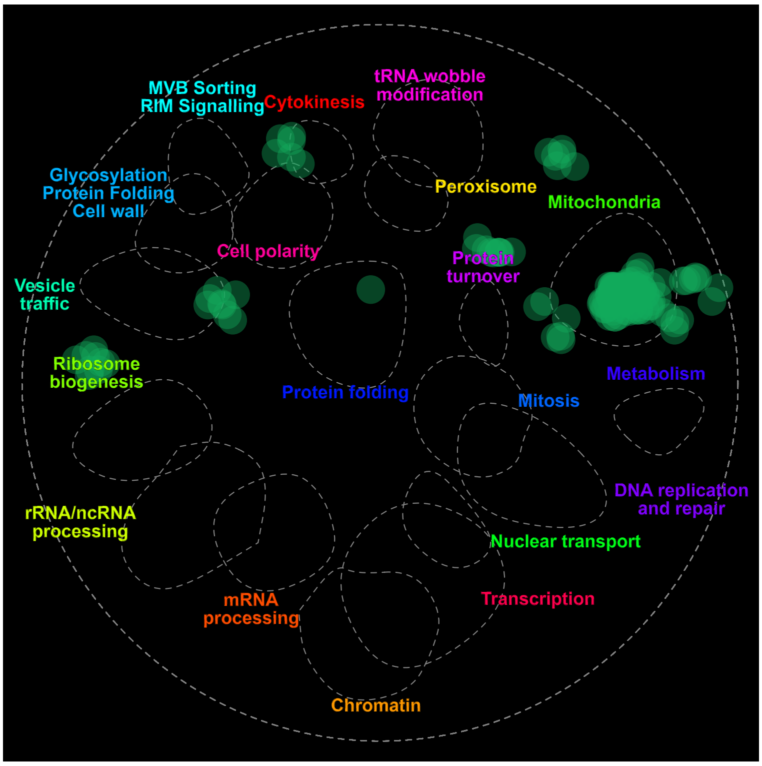


B

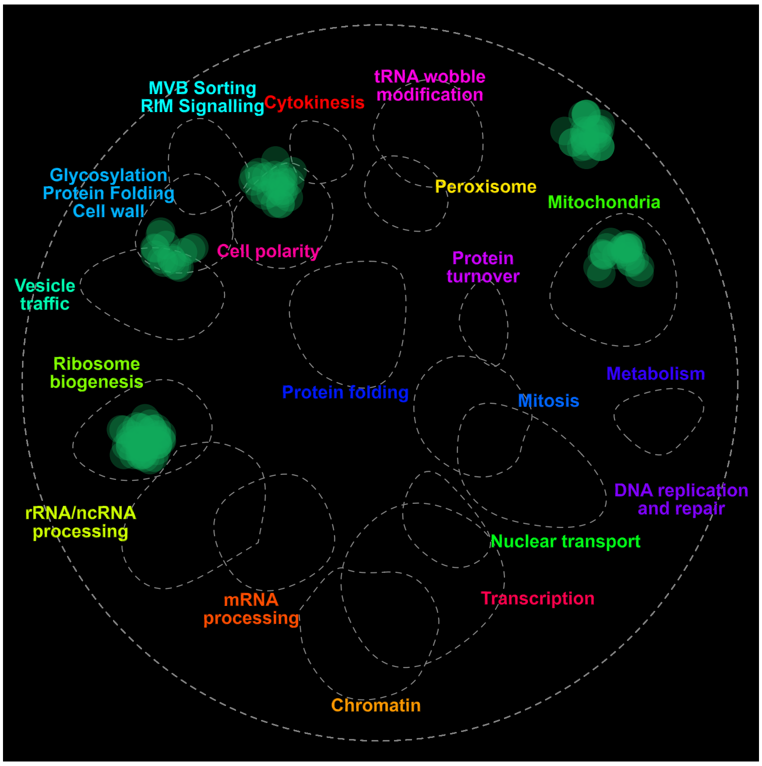


**Supplementary figure 1: SAFE analysis of synthetic lethal mutants and positive interactions with mip1∆.** Using the in-browser tool thecellmap.org (Usaj et al., 2017), non-dubious ORFs were submitted, with a significance cutoff of 0.05. **A:** Synthetic lethal mutants manually curated and derived from data. Two “dubious” ORFs were removed from the analysis, YNL170W and YDL062W. The remaining 17 genes were entered into thecellmap and revealed a significant enrichment in the category “Mitochondria” (p-value 2.34 E-14, fold change 4.12). **B:** Mutations leading to at least 50% better growth upon deletion of MIP1. 8 non-dubious ORF were submitted, excluding the dubious ORFs YIL012W, YOR309C and YPR063C. Significant enrichment was found in the category “Ribosome biogenesis” (p-value 2.29E-3, fold change 4.70) and “Glycosylation & protein folding” (p-value 2.29E-3, fold change 2.21). One strain deleted for the gene YPR022C predicted to encode for a transcription factor shows a synthetic rescue interaction.
