## Supplemental figure 2 for "Development of synthetic genetic arrays for studying interactions between the mitochondrial and nuclear genomes"

**A**

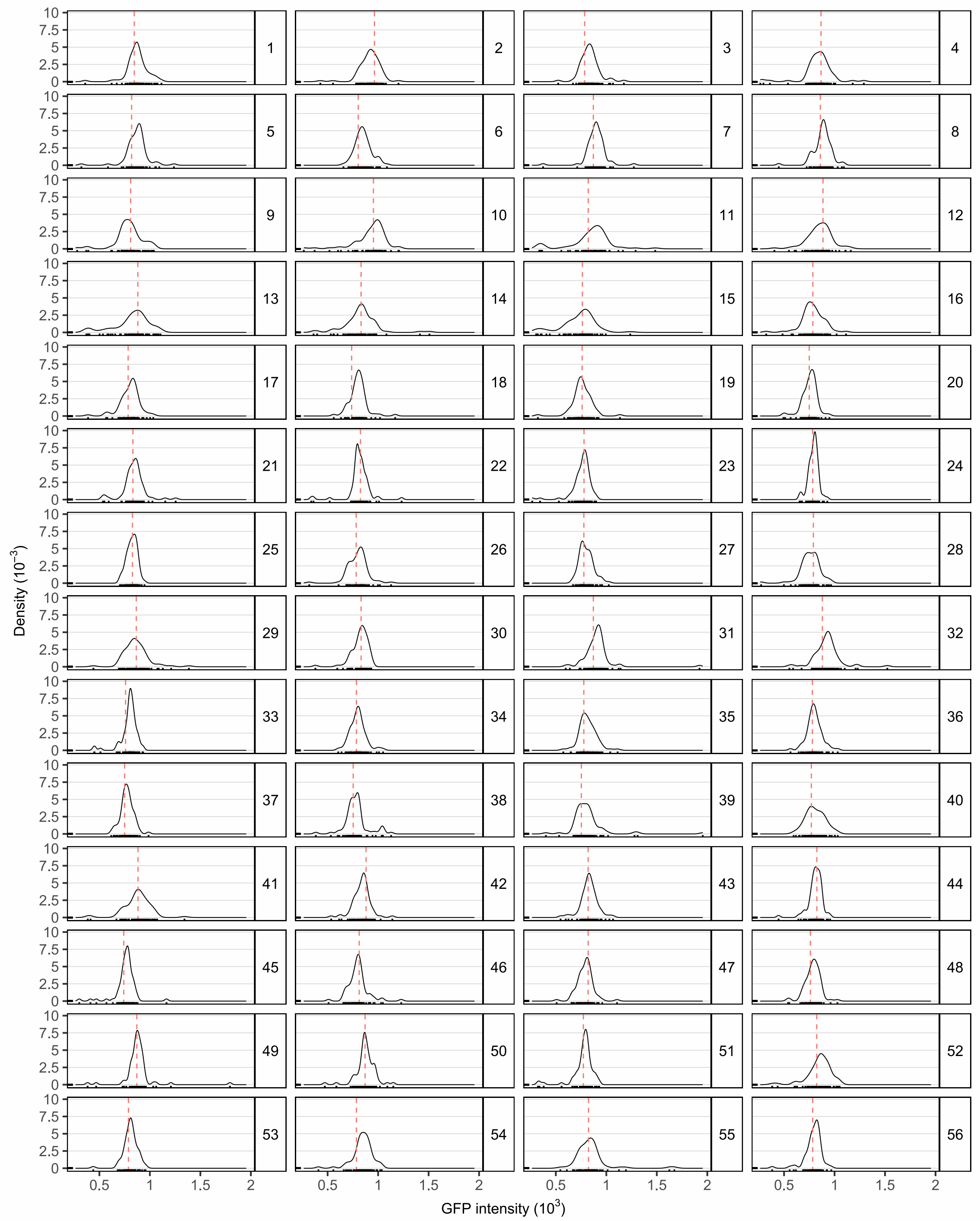
Density (10$^{-3}$)
GFP intensity (10$^{3}$)
0.5
1
1.5
2
0
2.5
5
7.5
10
1 2 3 4 5 6 7 8 9 10 11 12 13 14 15 16 17 18 19 20 21 22 23 24 25 26 27 28 29 30 31 32 33 34 35 36 37 38 39 40 41 42 43 44 45 46 47 48 49 50 51 52 53 54 55 56

**B**

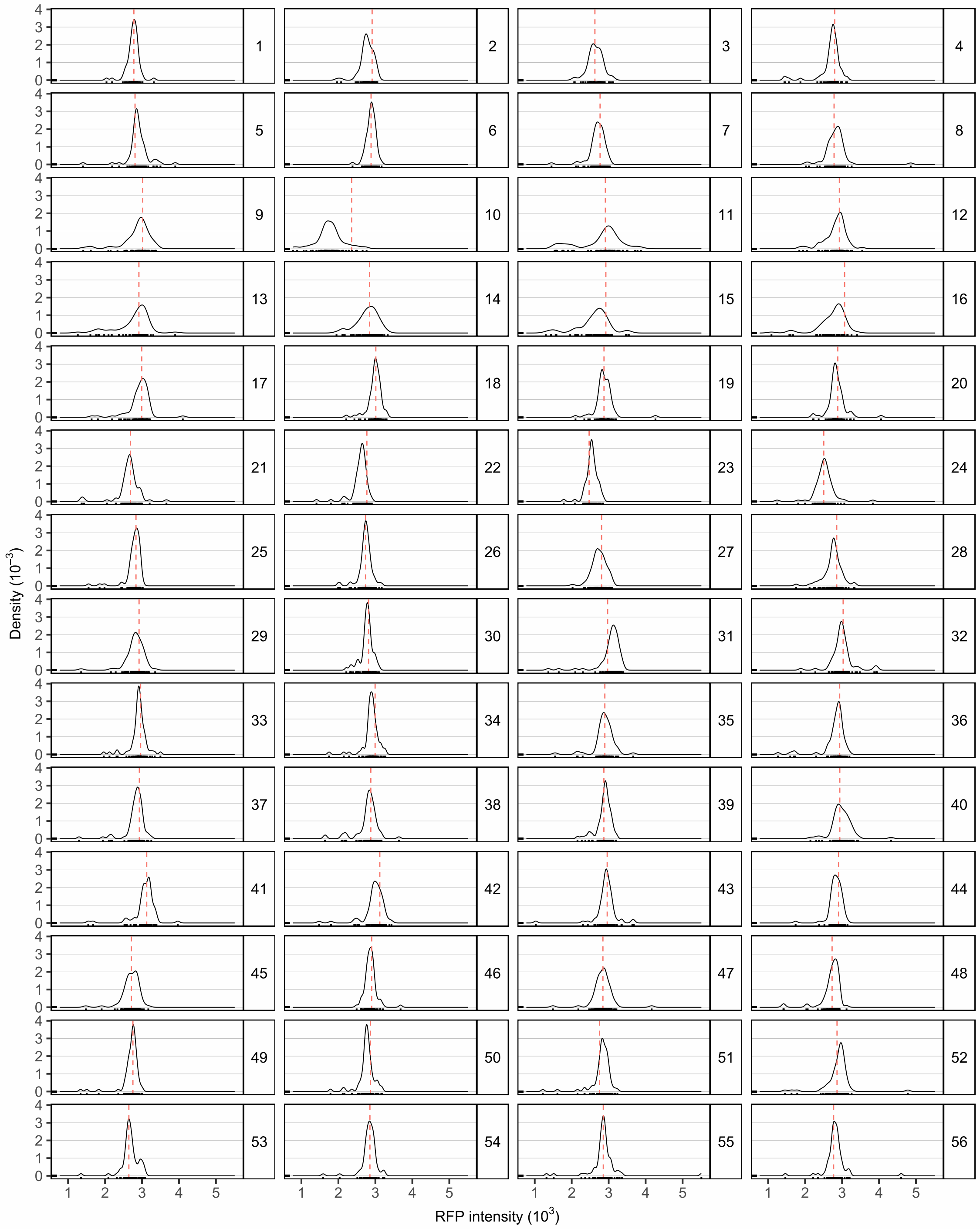
Density (10^-3)
RFP intensity (10^3)
1 2 3 4 5 6 7 8 9 10 11 12 13 14 15 16 17 18 19 20 21 22 23 24 25 26 27 28 29 30 31 32 33 34 35 36 37 38 39 40 41 42 43 44 45 46 47 48 49 50 51 52 53 54 55 56

**C**

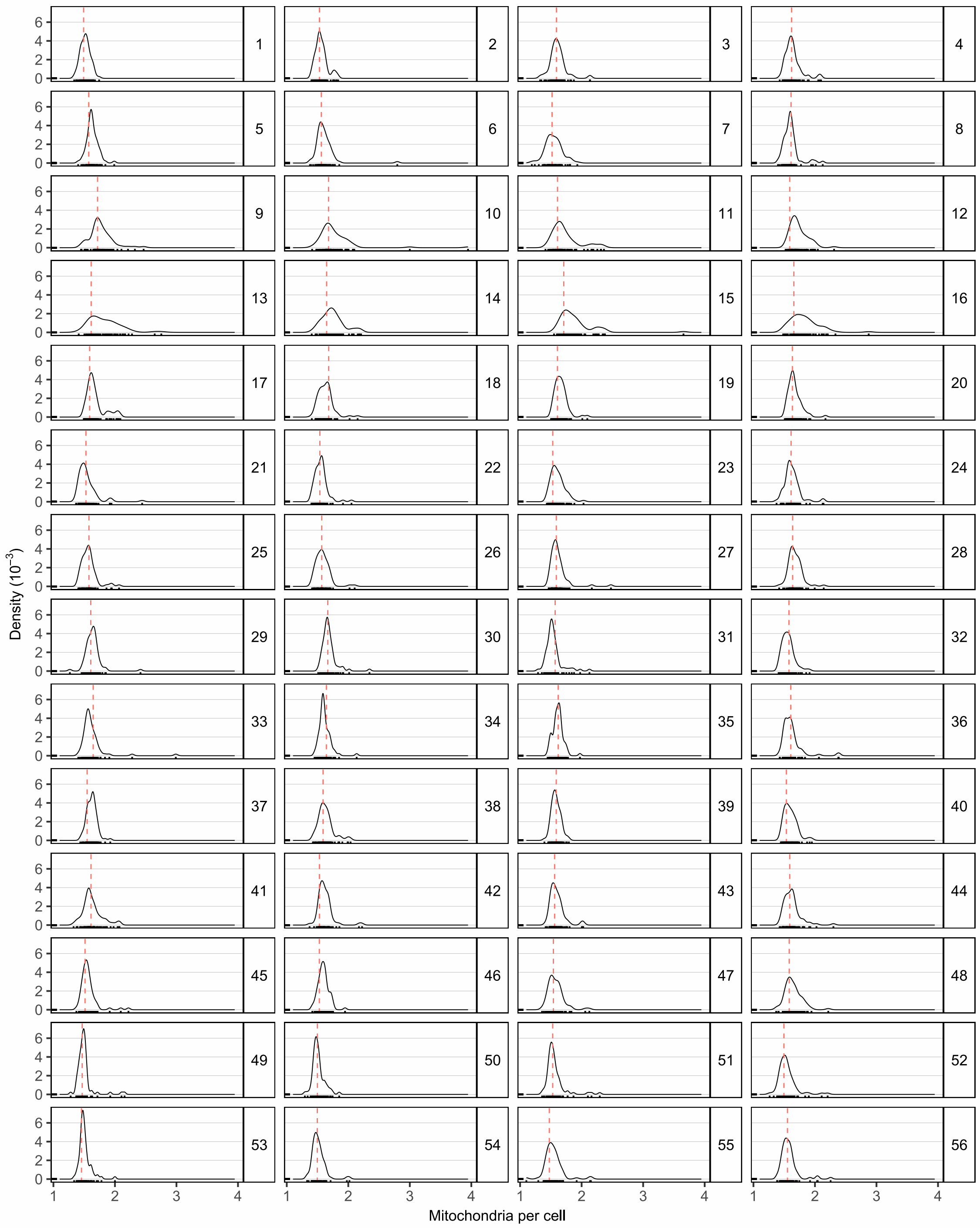
Density ($10^{-3}$)
Mitochondria per cell

**D**

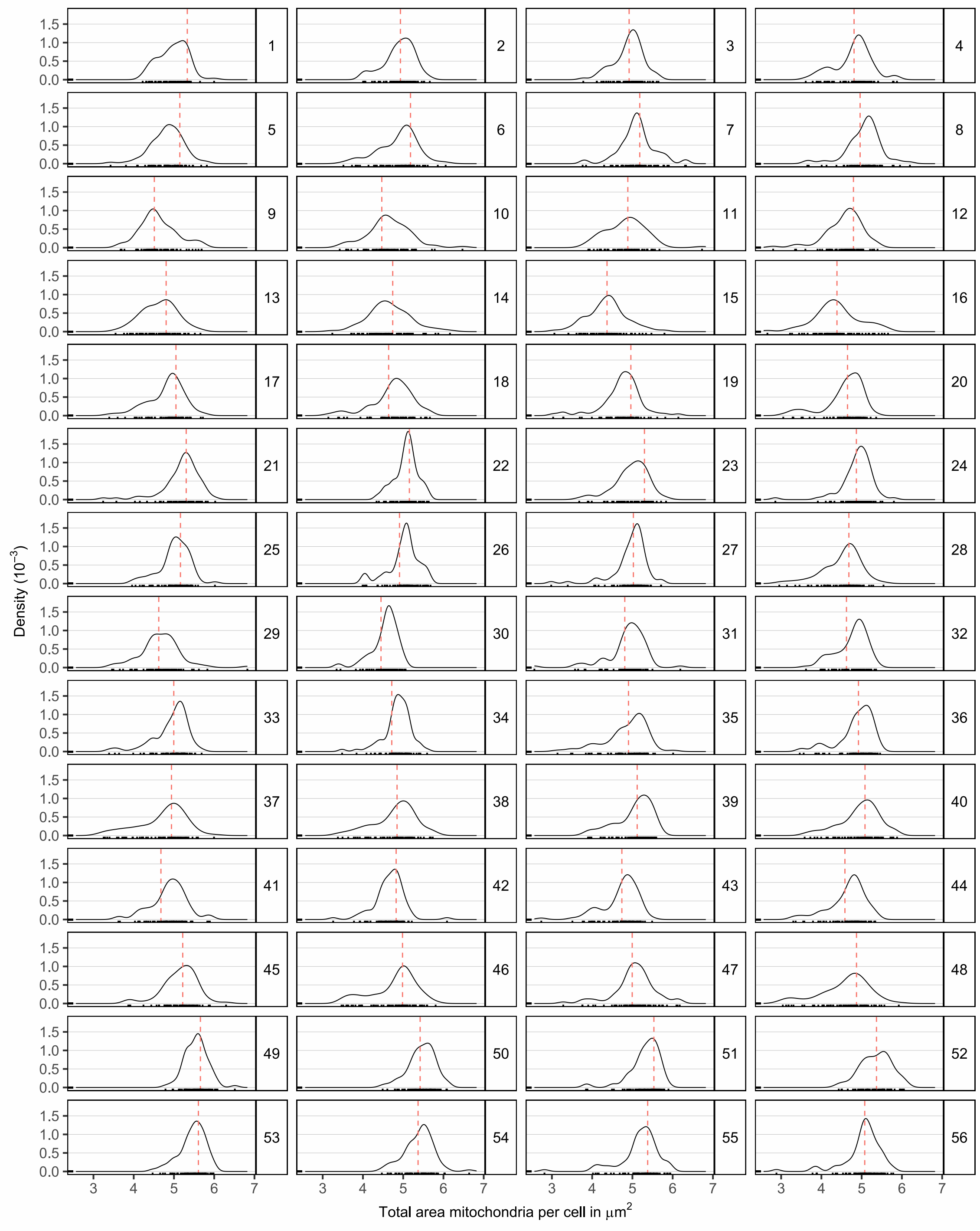

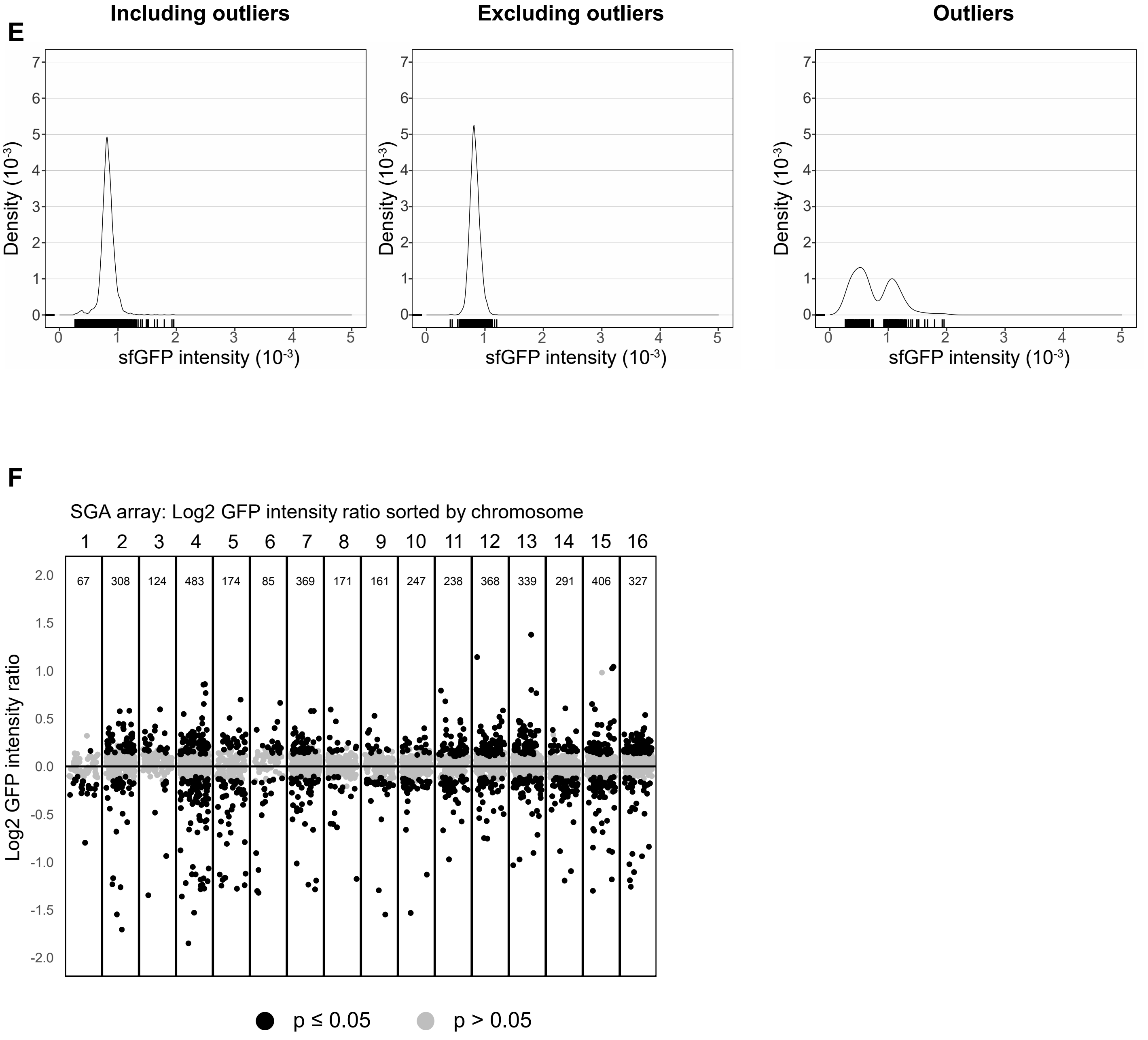

**Supplementary figure 2 A-D:** Plate-wise density distribution of phenotypes measured by GFP and RFP signal in single gene deletion strains. **A:** GFP intensity. **B:** RFP intensity of mitochondrial marker Tom70. **C:** mitochondrial count per cell. **D:** mitochondrial area per cell. Red dashed line represents weighed mean of control per plate, panel number represents plate number. **E:** Density distribution of GFP intensity summarized for all plates measured for single gene deletion test strains. Left: Including outlier, middle: excluding outliers, Right: distribution of outliers. **F:** Chromosome-wise plotting of -log2 of adjusted p-value, ordered after start location on chromosome. Black dots: significant adjusted p value ≤ 0.05, gray dots: non significant adjusted p-value. Panel numbers: chromosome number, n: number of observations per panel.
