## Supplemental figure 3 for "Development of synthetic genetic arrays for studying interactions between the mitochondrial and nuclear genomes"

**A**

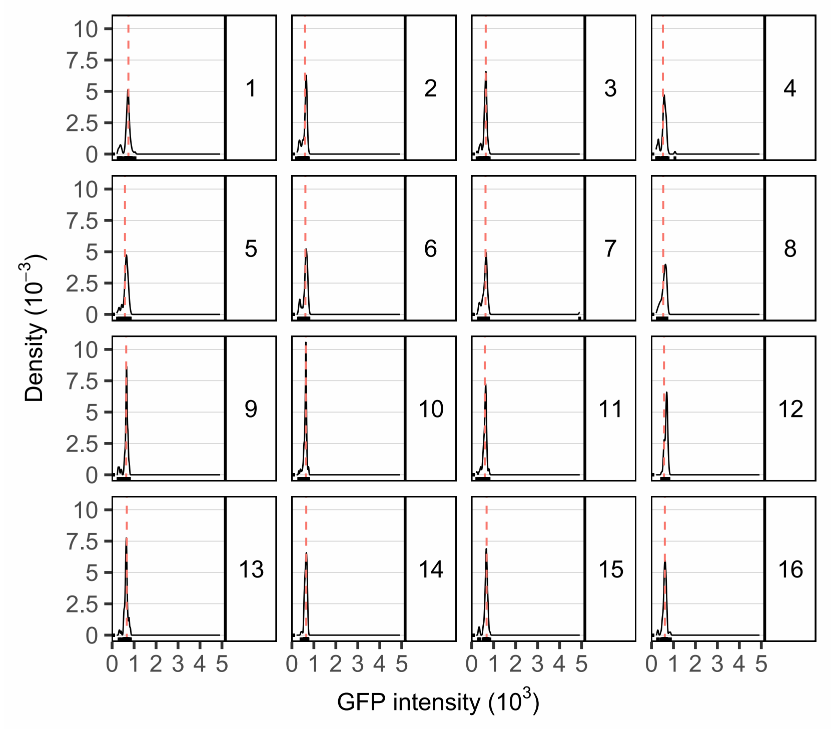

Density ($10^{-3}$)
GFP intensity ($10^3$)
10
7.5
5
2.5
0
0 1 2 3 4 5

**B**

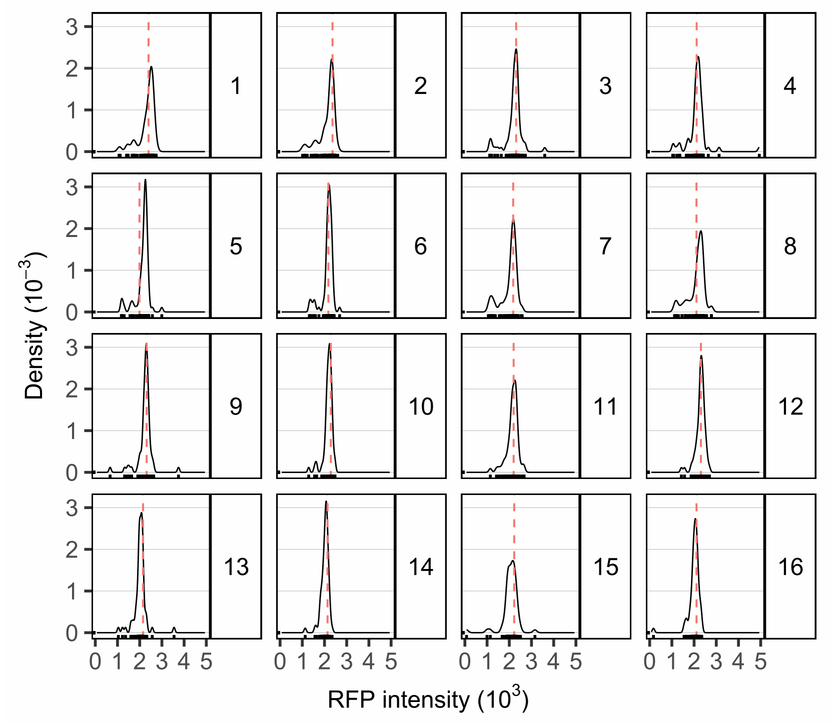

Density ($10^{-3}$)
RFP intensity ($10^3$)
3
2
1
0
0 1 2 3 4 5

**C**

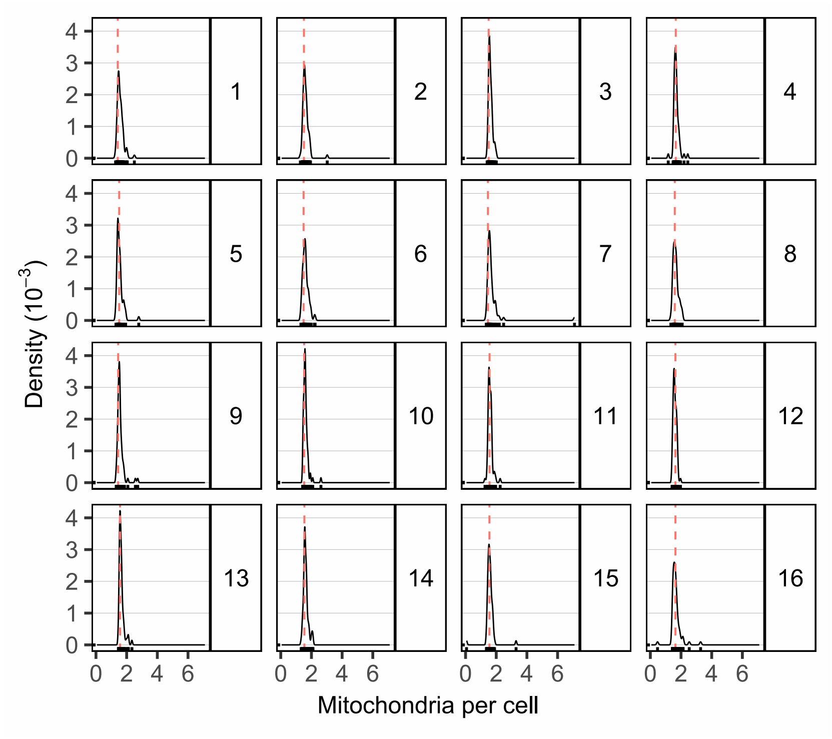

Density (10^-3)
Mitochondria per cell
0 2 4 6
0 1 2 3 4

**D**

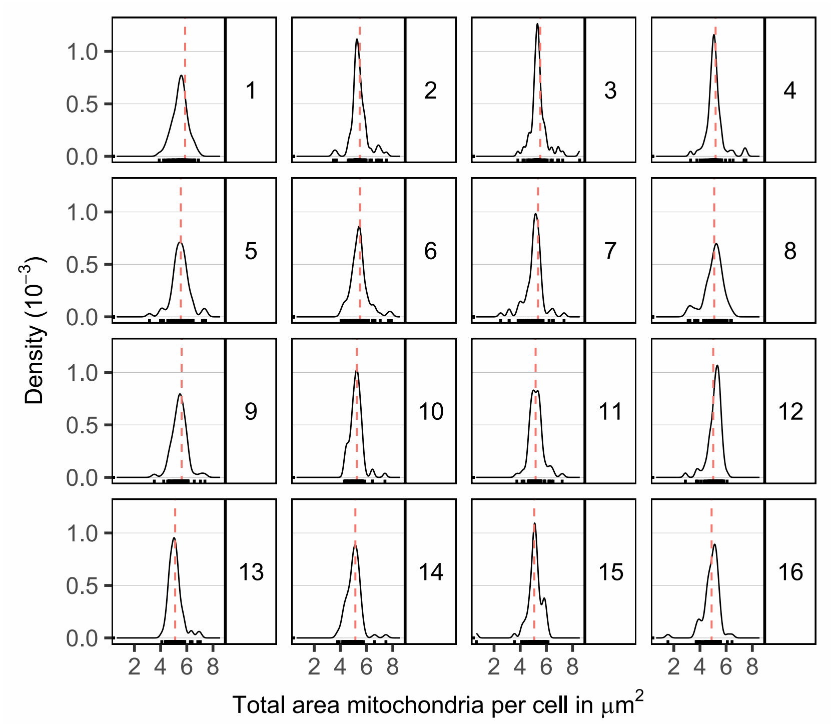

Density (10^-3)
Total area mitochondria per cell in μm^2
2 4 6 8
0.0 0.5 1.0

**E**

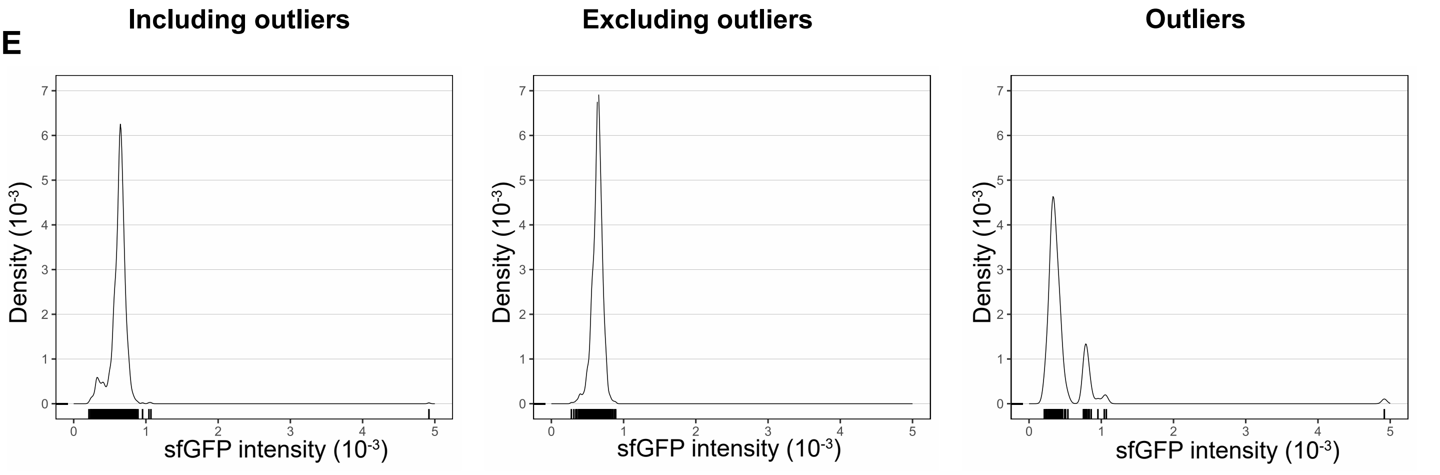
Including outliers
Excluding outliers
Outliers
Density (10^-3)
sfGFP intensity (10^-3)

**F**

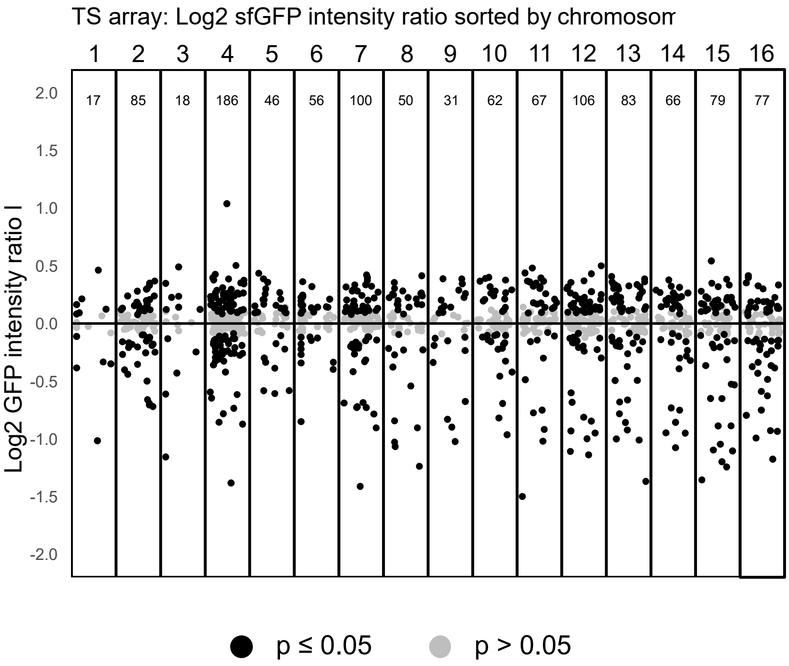
TS array: Log2 sfGFP intensity ratio sorted by chromosom
1 2 3 4 5 6 7 8 9 10 11 12 13 14 15 16
17 85 18 186 46 56 100 50 31 62 67 106 83 66 79 77
Log2 GFP intensity ratio I
p ≤ 0.05
p > 0.05

G

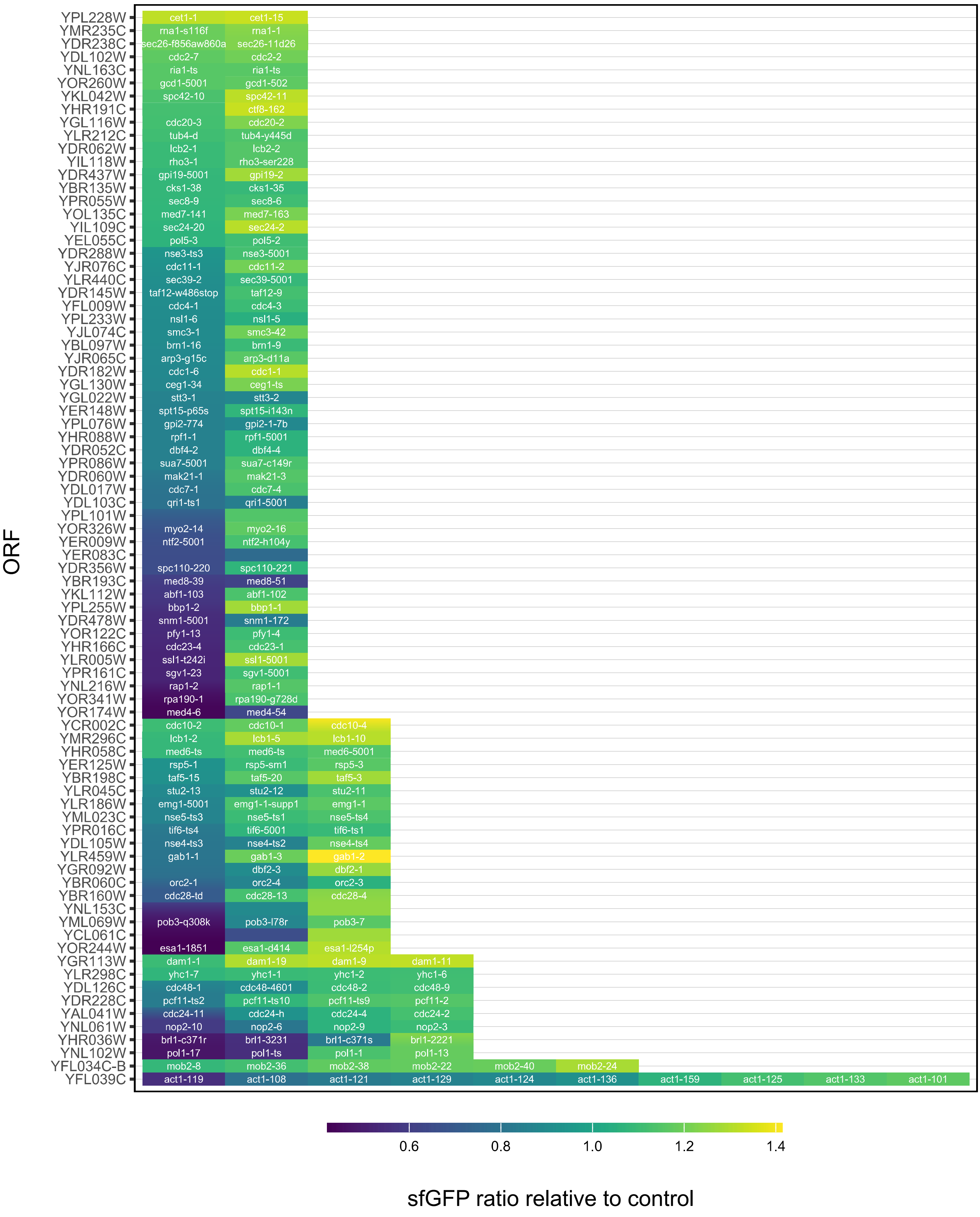

ORF
YPL228W cet1-1 cet1-15
YMR235C rna1-s116f rna1-1
YDR238C sec26-f856aw860a sec26-11d26
YDL102W cdc2-7 cdc2-2
YNL163C ria1-ts ria1-ts
YOR260W gcd1-5001 gcd1-502
YKL042W spc42-10 spc42-11
YHR191C ctf8-162
YGL116W cdc20-3 cdc20-2
YLR212C tub4-d tub4-y445d
YDR062W lcb2-1 lcb2-2
YIL118W rho3-1 rho3-ser228
YDR437W gpi19-5001 gpi19-2
YBR135W cks1-38 cks1-35
YPR055W sec8-9 sec8-6
YOL135C med7-141 med7-163
YIL109C sec24-20 sec24-2
YEL055C pol5-3 pol5-2
YDR288W nse3-ts3 nse3-5001
YJR076C cdc11-1 cdc11-2
YLR440C sec39-2 sec39-5001
YDR145W taf12-w486stop taf12-9
YFL009W cdc4-1 cdc4-3
YPL233W nsl1-6 nsl1-5
YJL074C smc3-1 smc3-42
YBL097W brn1-16 brn1-9
YJR065C arp3-g15c arp3-d11a
YDR182W cdc1-6 cdc1-1
YGL130W ceg1-34 ceg1-ts
YGL022W stt3-1 stt3-2
YER148W spt15-p65s spt15-i143n
YPL076W gpi2-774 gpi2-1-7b
YHR088W rpf1-1 rpf1-5001
YDR052C dbf4-2 dbf4-4
YPR086W sua7-5001 sua7-c149r
YDR060W mak21-1 mak21-3
YDL017W cdc7-1 cdc7-4
YDL103C qri1-ts1 qri1-5001
YPL101W
YOR326W myo2-14 myo2-16
YER009W ntf2-5001 ntf2-h104y
YER083C
YDR356W spc110-220 spc110-221
YBR193C med8-39 med8-51
YKL112W abf1-103 abf1-102
YPL255W bbp1-2 bbp1-1
YDR478W snm1-5001 snm1-172
YOR122C pfy1-13 pfy1-4
YHR166C cdc23-4 cdc23-1
YLR005W ssl1-t242i ssl1-5001
YPR161C sgv1-23 sgv1-5001
YNL216W rap1-2 rap1-1
YOR341W rpa190-1 rpa190-g728d
YOR174W med4-6 med4-54
YCR002C cdc10-2 cdc10-1 cdc10-4
YMR296C lcb1-2 lcb1-5 lcb1-10
YHR058C med6-ts med6-ts med6-5001
YER125W rsp5-1 rsp5-sm1 rsp5-3
YBR198C taf5-15 taf5-20 taf5-3
YLR045C stu2-13 stu2-12 stu2-11
YLR186W emg1-5001 emg1-1-supp1 emg1-1
YML023C nse5-ts3 nse5-ts1 nse5-ts4
YPR016C tif6-ts4 tif6-5001 tif6-ts1
YDL105W nse4-ts3 nse4-ts2 nse4-ts4
YLR459W gab1-1 gab1-3 gab1-2
YGR092W dbf2-3 dbf2-1
YBR060C orc2-1 orc2-4 orc2-3
YBR160W cdc28-td cdc28-13 cdc28-4
YNL153C
YML069W pob3-q308k pob3-l78r pob3-7
YCL061C
YOR244W esa1-1851 esa1-d414 esa1-l254p
YGR113W dam1-1 dam1-19 dam1-9 dam1-11
YLR298C yhc1-7 yhc1-1 yhc1-2 yhc1-6
YDL126C cdc48-1 cdc48-4601 cdc48-2 cdc48-9
YDR228C pcf11-ts2 pcf11-ts10 pcf11-ts9 pcf11-2
YAL041W cdc24-11 cdc24-h cdc24-4 cdc24-2
YNL061W nop2-10 nop2-6 nop2-9 nop2-3
YHR036W brl1-c371r brl1-3231 brl1-c371s brl1-2221
YNL102W pol1-17 pol1-ts pol1-1 pol1-13
YFL034C-B mob2-8 mob2-36 mob2-38 mob2-22 mob2-40 mob2-24
YFL039C act1-119 act1-108 act1-121 act1-129 act1-124 act1-136 act1-159 act1-125 act1-133 act1-101
0.6
0.8
1.0
1.2
1.4
sfGFP ratio relative to control

**Supplementary figure 3 A-D:** Plate-wise density distribution of phenotypes measured by GFP and RFP signal in temperature sensitive strains. **A:** GFP intensity. **B:** RFP intensity of mitochondrial marker Tom70. **C:** mitochondrial count per cell. **D:** mitochondrial area per cell. Red dashed line represents weighed mean of control per plate, panel number represents plate number. **E:** Density distribution of GFP intensity summarized for all plates measured for single gene deletion test strains. Left: Including outlier, middle: excluding outliers, Right: distribution of outliers. **F:** Chromosome-wise plotting of -log2 of adjusted p-value, ordered after start location on cchromosome. Black dots: significant adjusted p value $\leq$ 0.05, gray dots: non significant adjusted p-value. Panel numbers: chromosome number, n: number of observations per panel. **G:** Heatmap of change in GFP intensity ratio relative to control in genes with several allelic deletions within temperature sensitive array.
