## Supplemental table 3 for "Development of synthetic genetic arrays for studying interactions between the mitochondrial and nuclear genomes"

**Supplementary table 3:** Synthetic lethal double mutants selected by colony size and manual inspection of images and heatmap

| Identifier | Gene name | Score 1st run | Score 2nd run | Function Summary |
|---|---|---|---|---|
| YAL048C | GEM1 | -0.76479 | -0.62576 | Calcium-binding GTPase subunit of ERMES complex; involved in phospholipid homeostasis, mitochondrion-endoplasmic reticulum membrane tethering, mitochondrial organization and inheritance, and heme transport; localizes to outer mitochondrial membrane |
| YBR132C | AGP2 | -1.02236 | -- | Membrane protein that positively regulates the transport of (R)-carnitine and polyamines across membranes; localizes to plasma, vacuolar, and endoplasmic reticulum membranes |
| YCR028C-A | RIM1 | -0.71377 | -0.68013 | Single-stranded DNA binding protein involved in mitochondrial DNA replication; localizes to mitochondrial nucleoids |
| YCR071C | IMG2 | -0.49416 | -0.36315 | Protein constituent of large mitochondrial ribosomal subunit |
| YDL033C | SLM3 | -0.75003 | -0.80193 | Mitochondrial tRNA 2-thiouridylase involved in the thiolation of the wobble position uridine |
| YDL062W | Dubious | -0.80102 | -0.77823 | |
| YDL104C | QRI7 | -0.79976 | -0.86021 | Evolutionarily conserved mitochondrial N(6)-L-threonylcarbamoyladenine synthase involved in tRNA modification |
| YDR332W | IRC3 | -0.7804 | -0.80987 | Double-stranded DNA-dependent ATPase with a role in mitochondrial genome maintenance; localized to the mitochondrial matrix |
| YJR054W | KCH1 | -0.13737 | -0.54264 | Plasma membrane potassium transporter; part of the mating projection |

| | | | | |
|---|---|---|---|---|
| YLR114C | AVL9 | -0.78735 | -0.99723 | Protein involved in post-Golgi vesicle-mediated transport; localizes at cytoplasm, cellular bud neck and bud tip |
| YMR282C | AEP2 | -0.79129 | -0.5647 | Protein involved in translational initiation; localizes to mitochondria |
| YNL003C | PET8 | -0.7668 | -0.58202 | Mitochondrial S-adenosyl-L-methionine transmembrane transporter |
| YNL081C | SWS2 | -0.728 | -0.58011 | Component of the small subunit of the mitochondrial ribosome, which mediates translation in the mitochondrion; similar to bacterial small-subunit ribosomal proteins |
| YNL170W | Dubious | -0.91551 | -0.85681 | |
| YOL095C | HMI1 | -0.81321 | -0.67067 | 3'-5' DNA helicase involved in mitochondrial DNA metabolism; localizes to mitochondrial matrix |
| YOR033C | EXO1 | -1.04379 | -1.04153 | 5'-3' exonuclease and 5'-flap endonuclease involved in DNA double-strand break processing, gene conversion at mating-type locus, and mismatch repair; also involved in telomere maintenance and telomeric 3' overhang formation; localizes to the nucleus |
