## Supplemental table 4 for "Development of synthetic genetic arrays for studying interactions between the mitochondrial and nuclear genomes"

**Supplementary table 4:** Double mutants with positive interaction leading to increased growth over control. Mutants were chosen after calculating ratio of normalized colony size of mutant to control and candidates were chosen who showed at least 50% better growth compared to wildtype. Notes: * median, ** only rescue, present in both runs control shows zero growth, hence ratio couldn't be calculated normalized colony sizes for both runs are 0.5804 for the second and 0.41778 for the third run.

| Identifier | Gene name | Score 1st run | Score 2nd run | Function Summary |
|---|---|---|---|---|
| YEL031W | SPF1 | 0.25001 | 0.14122 | P-type ATPase involved in the homeostasis of calcium, manganese, and sterol; localizes to the endoplasmic reticulum membrane and the cis-Golgi network |
| YEL048C | TCA17 | 0.2965 | 0.40054 | Subunit of the TRAPPII complex, a multisubunit guanine nucleotide-exchange factor; involved in assembly of the TRAPPII complex and in early endosome to Golgi vesicle-mediated transport; localizes to the Golgi apparatus, and in high-throughput studies to the cytosol and clathrin-coated vesicles |
| YER089C | PTC2 | 0.35553 | 0.29352 | Serine/threonine phosphatase protein; dephosphorylates Hog1p, Ire1p, Cdk1p; negatively regulates endoplasmic reticulum (ER) unfolded protein response (UPR); localized to nucleus and cytoplasm |
| YFL036W | Dubious | 0.17615 | 0.15683 | Mitochondrial RNA polymerase; transcribes all mitochondrial genes and also synthesizes RNA primers for mitochondrial DNA replication; localizes to the mitochondrial matrix and associates with nucleoids |
| YGR157W | RPO41 | 0.35723 | 0.19782 | Phosphatidylethanolamine N-methyltransferase (PEMT) involved in phosphatidylcholine biosynthetic process; localizes to the endoplasmic reticulum |
| YIL012W | CHO2 | 0.28594 | 0.2755 | |
| YOL025W | LAG2 | 0.7227 | 0.67817 | Protein involved in negative regulation of protein ubiquitination |
| YOR309C | Dubious | 0.54221* | 0.4385 | |
| YOR312C | RPL20B | 0.44581 | 0.24971 | Subunit of the cytosolic large ribosomal subunit; involved in translation |
| YPR063C | Dubious | 0.57992 | 0.41744 | Protein whose biological role is unknown; localizes to the endoplasmic reticulum in a large-scale study |
| YPR022C** | SDD4 | 0.18957 | 0.34727 | Protein predicted by computational studies to act in transcription regulation |
